# Statistical modeling, estimation, and remediation of sample index hopping in multiplexed droplet-based single-cell RNA-seq data

**DOI:** 10.1101/617225

**Authors:** Rick Farouni, Haig Djambazian, Jiannis Ragoussis, Hamed S. Najafabadi

## Abstract

We introduce a probabilistic model for estimation of sample index-hopping rate in multiplexed droplet-based single-cell RNA sequencing data and for inference of the true sample of origin of the hopped reads. Across the datasets we analyzed, we estimate the sample index hopping probability to range between 0.003–0.009, a small number that counter-intuitively gives rise to a large fraction of ‘phantom molecules’ – as high as 85% in a given sample. We demonstrate that our model-based approach can correct for this artifact by accurately purging the majority of phantom molecules from the data. Code and reproducible analysis notebooks are available at https://github.com/csglab/phantom_purge.

**Structure:** Section 1 provides a concise summary of the paper. Section 2 provides a brief historical and technical overview of the phenomenon of sample index hopping and an explanation of related concepts. The three sections that follow describe the statistical modeling approach and correspond to the following three goals. (1) Building a generative model that probabilistically describes the phenomenon of sample index hopping of multiplexed sample reads (Section 3). (2) Estimating the index hopping rate from empirical experimental data (Section 4). (3) Correcting for the effects of sample index hopping through a principled probabilistic procedure that reassigns reads to their true sample of origin and discards predicted phantom molecules by optimally minimizing the false positive rate (Section 5). Next, Section 6 details the results of the analyses performed on empirical and experimental validation datasets. The Supplementary Notes consists of three sections: (1) Mathematical Derivations, (2) Overview of Computational Workflow, (3) Method’s Limitations.

## 1 Précis

Due to the increasing capacity of modern sequencing platforms, sample multiplexing, the pooling of barcoded DNA from multiple samples in the same lane of a high-throughput sequencer, is rapidly becoming the default option in single-cell RNA-seq (scRNA-seq) experiments. However, as several studies have recently shown^[3,10]^, multiplexing leads to incorrect sample assignment of a significant fraction of demultiplexed sequencing reads. Out of several mechanisms that can introduce sample index missassignment^[9]^, the presence of free-floating indexing primers that attach to the pooled cDNA fragments just before the exclusion amplification step in patterned sequencing flowcells has been shown to be the main culprit^[6]^. This phenomenon is known as *sample index hopping* and results in a data cross-contamination artifact that takes the form of *phantom molecules*, molecules that exist only in the data by virtue of read misassignment (Fig. 1a). The presence of phantom molecules in droplet-based scRNA-seq data should be a cause of great concern since they can introduce both phantom cells and artificial differentially expressed genes in downstream analyses. Importantly, it is conceivable that even when the index-hopping rate is very small, the fraction of phantom molecules can still be high due to the distributional properties of sequencing reads across samples (Fig. 1b).

**Figure 1:**
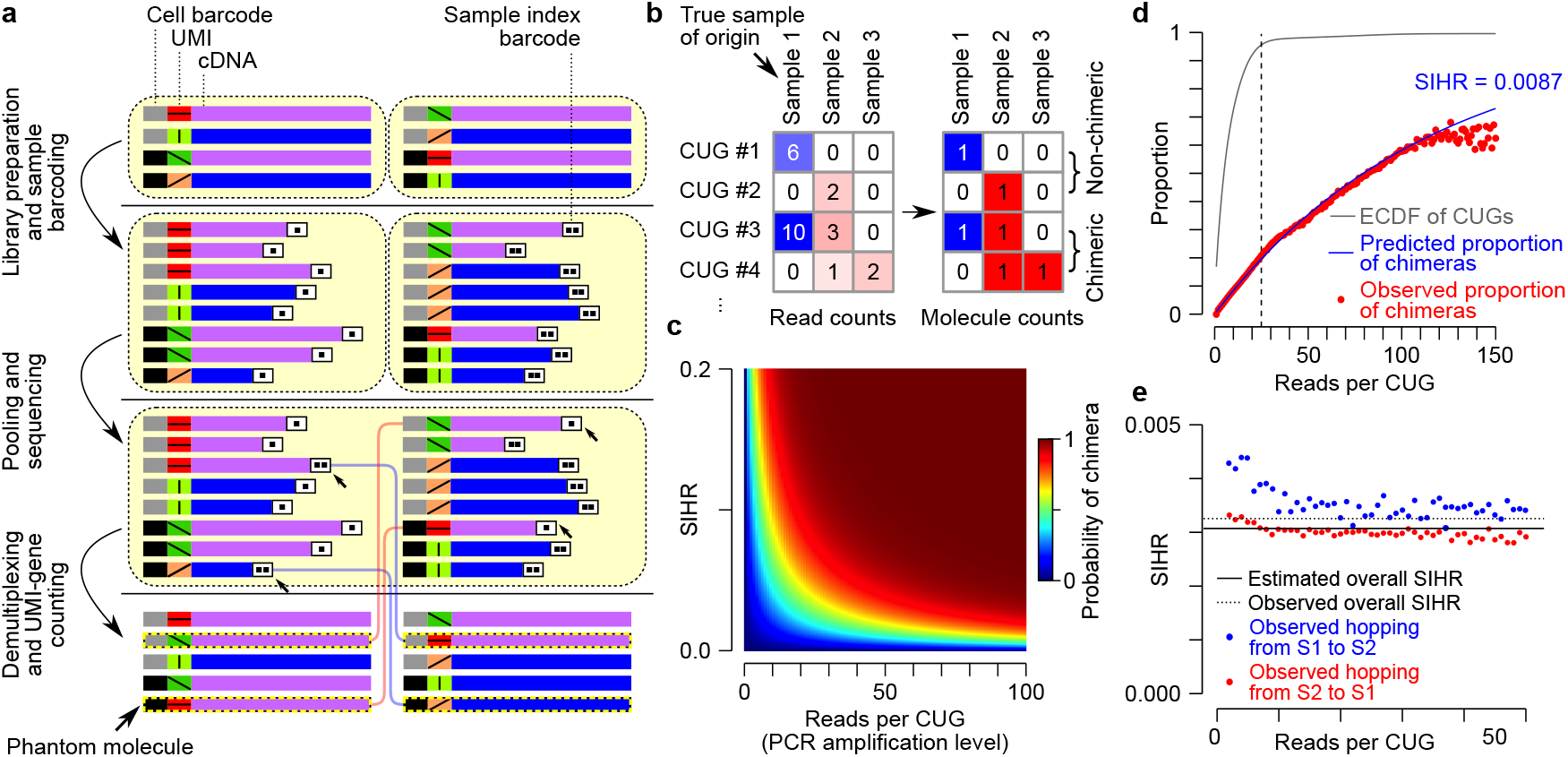
(**a**) A schematic representation of of sample index hopping. (**b**) Example count matrices showing hopped reads (left) and the resulting molecule count matrix (right). Blue and red depict true sample of origin and the hopped sample, respectively, with the color intensity showing relative counts. (**c**) A heatmap depicting the probability of observing a chimera as a function of hopping rate (SIHR) and PCR amplification level *r*. (**d**) Model fit showing predicted mean function overlayed over the observed proportion of chimeric observations for a HiSeq 4000 multiplexed dataset^[1]^. (**e**) Validation data showing the ground-truth proportion of hopped reads by sample conditional on the PCR duplication level alongside the ground-truth marginal mean proportion of hopped reads and the model estimated sample index hopping rate.

Despite recent attempts to computationally estimate the rate of sample index hopping in plate-based scRNA-seq data^[4,7]^, no statistical model of index hopping for droplet-based scRNA-seq data has yet been proposed. Consequently, current computational methods can neither accurately estimate the underlying rate of index hopping nor adequately remove the resulting phantom molecules in droplet-based scRNA-seq data. This has been a challenging problem since droplet-based libraries are tagged with a single sample index rather than a unique combinatorial pair of sample indices such as those used in plate-based approaches. As a solution to this problem, we here propose a statistical framework that provides (i) a generative probabilistic model that formalizes in a mathematically rigorous manner the phenomenon of index hopping, (ii) a statistical approach for inferring the sample index hopping rate (SIHR) in droplet-based scRNA-seq data at the level of individual reads, (iii) a non-heuristic, model-based approach for inferring the true sample of origin of hopped sequencing reads, and (iv) a data decontamination procedure for purging phantom molecules that optimally minimizes the false positive rate of molecule re-assignments.

The generative probabilistic model we propose starts with the observation that each cDNA fragment, in addition to its sample barcode index, has a cell barcode and a unique molecular identifier (UMI), and maps to a specific gene. As has been suggested previously^[4]^, we make the assumption that any particular cell-UMI-gene combination (hereafter referred to as CUG) is so unlikely that it cannot arise independently in any two different samples. Accordingly, each CUG would represent one unique molecule and all sequencing reads with the same combination would correspond to PCR amplification products of that original molecule. A second assumption we make is that the probability of index hopping is the same for all reads, regardless of the source or target sample of the read (**Methods 3.3**). We were able to validate both assumptions (Fig. 8 and **Supplementary Notes**) by analyzing data from an experiment in which two 10X Genomics scRNA-seq sample libraries were sequenced in two conditions. In the first condition, two sample libraries were multiplexed on the same lane of HiSeq 4000. In the second condition, the same libraries were sequenced separately on two lanes of HiSeq 4000 (this non-multiplexed condition provides a ground truth for the true sample of origin of each CUG; see **Supplementary Notes 5**). The validation dataset has been deposited on the open-access data repository Zenodo (https://doi.org/10.5281/zenodo.3267922)

Building on these two assumptions, we derived a mixture-of-multinomials model (**Methods 3.4**) whose likelihood is governed by a single "index hopping" parameter that determines the observed distribution of read counts across samples at each PCR amplification level. We used this model to further derive a closed-form expression of the probability distribution of *chimeric* observations, namely CUGs whose corresponding reads are assigned to multiple samples (Fig. 1c and **Methods 4.1**). We were then able to estimate the index hopping rate by fitting the resulting generalized linear model to the empirically observed distribution of chimeric CUGs across PCR amplification levels (**Methods 4.4**). We observed close agreement between the observed and fitted non-chimeric CUGs across multiple datasets, including two eight-sample HiSeq datasets from mouse epithelial cells^[1,4]^ and two 16-sample NovaSeq 6000 datasets from Tabula Muris^[2]^ (Fig. 1d and Fig. 5). We also observed excellent agreement between our SIHR estimate and the empirical estimate for the dataset with known ground truth (Fig. 1e). Overall, we found that SIHR ranges between 0.3% and 0.9% in the datasets we analyzed (Table 4).

Furthermore, the proposed framework also allows the calculation of the posterior distribution of the true sample of origin for each CUG, given its read counts across samples, the SIHR, and the molecular proportions complexity profile of the samples (Fig. 2a and **Methods 3.5**). It then assigns each CUG to the most likely sample of origin, and further decontaminates the data by removing assignments that have low posterior probability. We have devised an approach for estimating the number of false positives (FP) and false negatives (FN) for distinguishing true molecules from phantom molecules after decontamination at different posterior probability cutoffs (Fig. 6), enabling the selection of a cutoff based on a user-specified marginal trade-off ratio (TOR) that represents the number of real molecules one is willing to discard in order to correctly purge one extra phantom molecule (Fig. 7 and **Methods 5.3**). These model-based FP and FN estimates are in excellent agreement with empirical estimates based on ground truth (Fig. 2b). We observed that we can achieve a sensitivity of 0.999 (down from 1 in the original non-purged data) and specificity of 0.979 (up from 0 in the non-purged data) in distinguishing true molecules from phantom molecules across all samples. Furthermore, our model-based purging of phantom molecules substantially outperforms, both on validated and simulated data, a previous heuristic approach^[4]^ that is based on retaining CUGs with a certain minimum fraction of reads (MRF) assigned to one sample (Fig. 2b, Fig. 10, and Tables 5–11).

**Figure 2:**
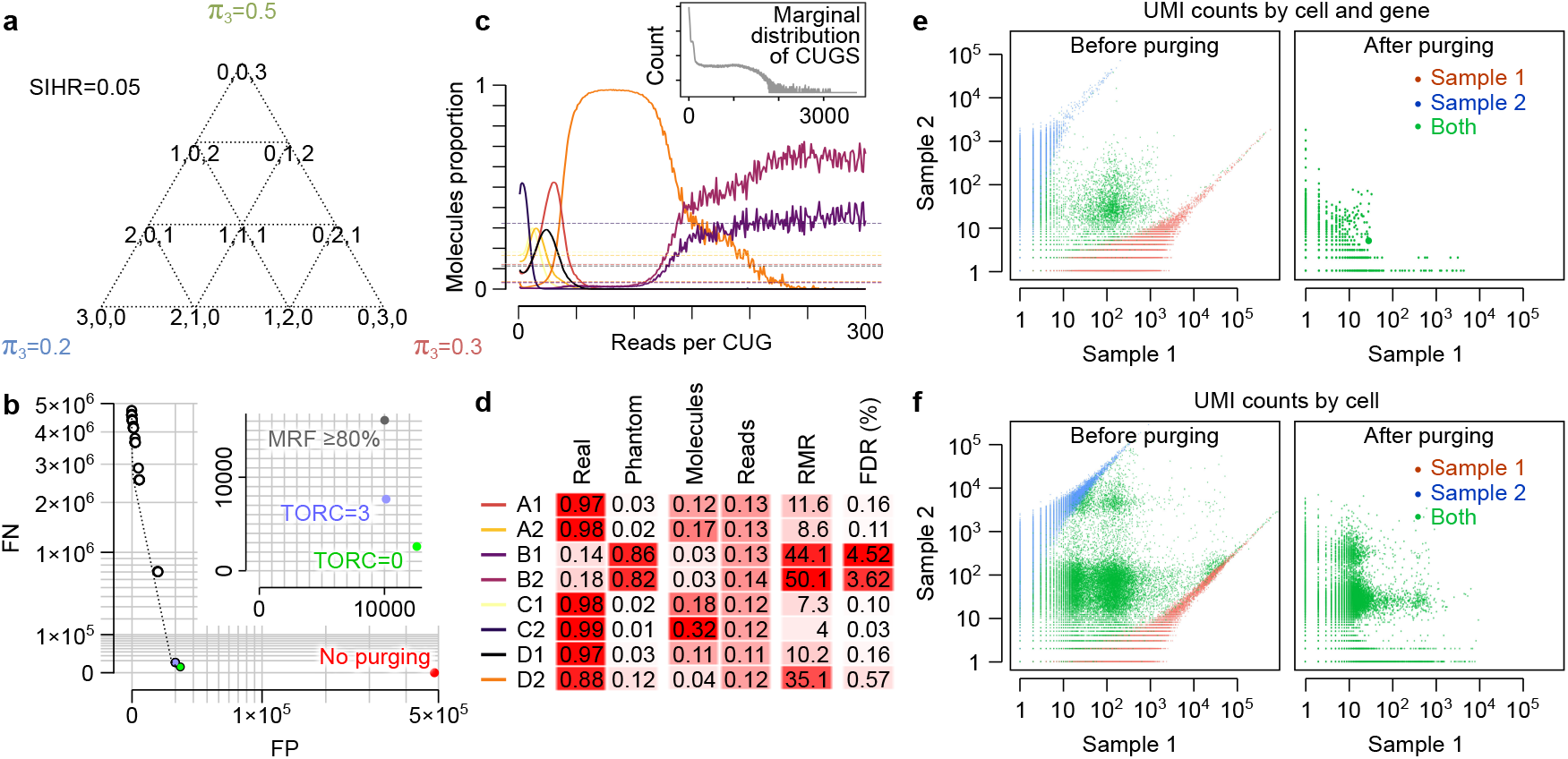
(**a**) The simplex containing all 10 possible read count outcomes at PCR amplification level (*r* = 3) and for the case of 3 samples. The pie charts depict the posterior probabilities of the true sample of origin for each outcome. *π*_3_ represents the proportion of molecules with *r* = 3 that originate from each sample in this toy example. (**b**) Effect of purging and discarding on the number of false positive and false negative counts. Points closer to the origin are more optimal. FP: false positive; FN: false negative. (**c**) The molecular library complexity profile for eight samples in a previously published HiSeq 4000 dataset^[1]^. (**d**) The proportion of real and phantom molecules in each of the eight samples, the fraction of molecules and mapped reads in the entire dataset that belong to each sample, the read-to-molecule ratio (RMR) per sample, and the corresponding FDR statistics (i.e. the within-sample proportion of molecules that we miss-classify as real after purging). (**e**) The effect of purging on gene expression profiling in the validation dataset with known ground truth. Each dot represents one gene expression measurement in one cell. Dots are colored based on cell-sample assignment in the ground truth, with red and blue representing cell barcodes that are found only in Sample 1 or Sample 2, respectively. Green represents cell barcodes that are found in both samples (barcode collision). Note that non-zero Sample 1 UMI counts for blue dots and non-zero Sample 2 UMI counts for red dots represent phantom molecules. (**f**) Same as panel (e), but with the counts aggregated per cell (each dots represents one cell). Blue dots with high counts in Sample 1 and red dots with high counts in Sample 2 represent phantom cells. Green dots represent potential transcriptome mixing.

Surprisingly, we found that the proportion of phantom molecules (PPM) varies widely depending on the *molecular proportions complexity profile* (Fig. 2c and Fig. 4) of the samples, and can reach as high as 86% in low-complexity samples (i.e. samples in which a relatively small number of unique molecules contribute to the majority of sequencing reads, Fig. 2d and Tables 7–8). We believe that high PPM values might in fact be common in multiplexed datasets that contain both low- and high-complexity samples, since hopping of even a small fraction of reads from high-to low-complexity samples can create more phantom molecules than the unique real molecules native to the low-complexity sample. Overall, these results indicate that even a small sample index hopping rate can be a substantial confounding factor in multiplexed scRNA-seq experiments by overwhelming the data with phantom molecules. However, the effects of index hopping on the integrity of the data, whether it leads to mixing of transcriptomes or creation of phantom cells (Fig. 9), or the miss-classification of RNA-containing cells vs. empty droplets (Tables 9–10), can be almost completely remedied by model-based purging of phantom molecules (Fig. 2e-f). The code and reproducible analysis notebooks for our approach are available at https://github.com/csglab/phantom_purg.

## 2 Background

This section describes the phenomenon of sample index hopping and provides a brief overview of existing published research that aims to quantify its rate and to correct for its data corrupting effects. The section also highlight the limitations of existing approaches and argues that the issue of index hopping is a significant problem in the field, one that no satisfactory solution for yet exists.

### 2.1 Overview of Index Hopping on Illumina’s Sequencers

#### Patterned flowcells

Ever since the emergence of next-generation sequencing technologies in 2006, the speed and throughput at which we can perform whole-genome DNA and RNA sequencing have been steadily increasing. For example, in recent years Illumina has introduced a family of sequencing platforms (i.e. the HiSeq 4000 and NovaSeq 6000) characterized by substantial increased capacity, faster run times, and lower sequencing cost. The significant boost in sequencing throughput^1^ first came about in 2015 when Illumina introduced the patterned flow cell technology on the HiSeq 4000 sequencers. In particular, there were two key innovations that led to the significantly improved performance. (1) A flow cell surface design that optimizes the prearranged spacing of nanowells, thus allowing the accurate imaging of billions of clusters of amplified cDNA fragments. (2) *The Exclusion Amplification* (ExAmp) chemistry that instantaneously amplifies a single cDNA fragment, effectively excluding and preventing other cDNA fragments from forming a cluster within the same nanowell. In 2017, Illumina launched the NovaSeq 6000 instrument with an updated flow cell design that further decreased the spacing between the nanowells, thus significantly increasing the density of generated clusters and, consequently, the amount of data that can be generated. Currently, a total of 2.5B single reads can be generated on each of the four lanes that comprise a single NovaSeq S4 flow cell, compared to 350M reads on each of eights lanes of HiSeq 4000.

#### Sample multiplex sequencing

However, given that for most sequencing studies, the number of reads required to achieve an adequate transcriptome coverage tends to be in the tens of millions, it then becomes necessary to resort to *sample multiplex sequencing* strategies to fully utilize the massive capacity and cost efficiency that the patterned flow cell with ExAmp chemistry technology can bring about. The process of sample multiplexing can be briefly summarized with the following few steps. First, a *sample barcode index* is added to each cDNA fragment during library preparation (to one end in *single-indexing* or to both ends in *dual-indexing*). Then, cDNA fragments from multiple samples are subsequently pooled together in the same lane to be cluster-amplified and sequenced. Finally, the generated sequenced reads are demultiplexed into their respective source samples using the sample barcode indices. Note that with current technology, in a single run of droplet-based RNA-seq experiment, a maximum of a 384 samples can be multiplexed (Chromium i7 Multiplex Kit) using a 96-plex on each of the four lanes of a NovaSeq S4 flow cell. In what follows, we define a *sequencing library* as a collection of index-barcoded and PCR-amplified cDNA fragments which are purified from the mRNA of a particular sample tissue

#### Sample read misassignment

Unfortunately, sample multiplexing can cause a significant percentage of the demultiplexed sequenced reads to be misassigned to an incorrect sample barcode. Although sample read misassignments can arise due to several factors^[9]^, one specific mechanism termed *sample index hopping* is the primary cause of read misassignments in patterned flow cells. Index hopping is believed to result from the presence of free-floating indexing primers that attach to the pooled cDNA fragments just before the exclusion amplification step that generates clusters on the flow cell. In addition, if during library preparation, free adapters or primers are not properly removed, the resulting purified library would show higher levels of index hopping that increases linearly with the molar concentration of free adaptors relative to DNA input^[6]^.

### 2.2 Discovery and Quantification of Index Hopping

The problem of index hopping was first identified as early as December 2016 in a blog post reporting read sample misassignment on HiSeq 4000 and HiSeqX platforms^[5]^. Soon afterwards, Illumina^[6]^ released a white paper acknowledging that index switching does indeed occur and tends to be higher in machines that use a patterned flow cell, but maintaining that the phenomenon affects only <2 % of reads. However, around the same time, Sinha *et al.*^[10]^ reported that 5-10% of sequencing reads were incorrectly assigned a sample index in a multiplexed pool of plate-based scRNA-seq samples. More recently, using two plate-based scRNA-seq datasets, Griffiths *et al.*^[4]^ provided a lower estimate of the index swapping on the HiSeq 4000 at approximately 2.5%. Another study^[11]^ conducted in the context of exome sequencing also reported a contamination (index hopping rate) ranging from 2% to 8%. Using unique antigen receptor expression,^[12]^ estimated the index hopping rate in plate-based single cell RNA-seq data (i.e. spread-of-signal across wells) to be approximately 3.9%. They also failed to detect any evidence that sample barcode indices vary in their proneness to undergo index swapping than others. With the aim of reconciling the conflicting hopping rate estimates that had been reported, Costello *et al.*^[3]^ used a non-redundant dual indexing adapters developed in-house and performed an exhaustive study across multiple libraries (whole genomes, exome, and stranded RNA) and sequencer models (HiSeq 4000, NovaSeq 6000 and HiSeqX) to determine the rate of index hopping. They observed a rate of 0.2% to 6% in all sequencing runs. More importantly, they showed that even in bulk RNA-seq libraries where the indexing hopping rate was as low as 0.32%, spurious results in downstream analysis can be generated. This finding mirrors the conclusions reached by the other five papers referenced in this paragraph including Illumina’s white paper.

### 2.3 Impact and Significance of Index Hopping

Given the large amount of data that are being increasingly generated using multiplexed sequencing, index hopping should be a great cause of concern due to potential signal artifacts and spurious results that it can generate. Here we list the three main unwanted consequences in the context of scRNA-seq sequencing experiments that sample index hopping can bring about through the introduction of phantom molecules.

1. *Mixing of transcriptomes*: when a given cell-barcode is observed in both the donor and target samples, index hopping would result in phantom molecules being introduced in the cell with the corresponding cell-barcode in the target sample. When this happens, the rates of false positives and even false negatives can increase in downstream statistical analyses such as differential gene expression.
2. *Phantom cells*: when a given cell-barcode is observed only in the donor sample, but not the target, phantom molecules would also lead to phantom cells emerging in the target sample.
3. *Cell-barcode miss-classification*: an abundance of phantom molecules associated with a given cell-barcode would lead cell-barcoding algorithms (used to computationally determine which cell-barcodes originate from proper cells) to incorrectly classify an empty droplet as a cell and conversely to classify a proper cell as an empty droplet.

### 2.4 Mitigating the Effects of Sample Index Hopping

#### 2.4.1 Experimental Strategies

Given that index hopping would occur in all sequencing platforms that uses patterned flow cells, one can resort to any of the following three experimental strategies to mitigate sample index hopping from affecting the integrity of experimental data.

1. *One sample per lane*. By avoiding multiplexing altogether and running one sample per lane, one can confine the sample indices in a given lane to be for one given sample only, thus minimizing the risk of index hopping from occurring in the first place. However, sample read misassignments can still occur due to other causes.
2. *Unique-at-both-ends dual indexing library*. By utilizing two unique barcodes for each sample, sample misassignment would occur only when both sample indices hop. Given the low probability that such an event would occur, index hopping can be reduced and computationally mitigated by discarding reads with unexpected combinations post-hoc.
3. *Post-library prep treatment*. By using Illumina’s Free Adapter Blocking Reagent, the 3^*1*^ ends of the free adapters become blocked preventing their extension and thus reducing the rate of index hopping^[6]^.

##### Limitations of Experimental Strategies

Although running one sample per lane can be feasible on HiSeq platforms for some single cell RNA-seq libraries, it would be financially prohibitive on newer higher capacity NovaSeq platforms. As for unique-at-both-ends indexing, the strategy is currently incompatible with single index droplet-based protocols such as the widely used 10x Genomics single cell protocol. Lastly, the blocking reagent Illumina provides is only compatible with a few bulk DNA and RNA protocols, but not for single cell protocols. Whether it can be incorporated in single cell droplet-based protocol is yet to be determined.

#### 2.4.2 Computational Strategies

Given the limitations of existing experimental strategies to eliminate index hopping, the urgency to develop computational strategies instead has only intensified since the phenomenon was first discovered. Indeed, the first attempt to computationally correct for index hopping in multiplexed sequencing libraries was only recently published^[7]^. However, the linear regression method the authors propose is applicable only to plate-based scRNA-seq samples, where a unique combination of a pair of sample barcodes in each well determines the identity of cells and can give rise to "crosshair" pattern in the data when index hopping occurs. It is this pattern that allows the estimation of fraction of hopped reads and upon which their proposed approach relies. Another limitation of their approach is that it uses only a subset of the data corresponding to genes whose specific cell expression is above a given threshold. However, Griffiths *et al.*^[4]^ developed a more accurate method for plate-based scRNA-seq that makes fewer assumptions and that uses data from all the genes. In the same paper, the authors also proposed a heuristic computational strategy for droplet-based scRNA-seq 10x Genomics experiments that excludes hopped reads without removing the corresponding cell libraries, but did not attempt to propose a modeling framework for estimating sample index hopping in droplet-based data since unlike plate-based assays, droplet-based assays do not use a unique pair of sample barcodes but rather use one mate of the paired read for quantification and a second mate to carry the UMI and cell barcode tags, thus rendering the "crosshair" pattern approach inapplicable.

##### Limitations of existing computational strategies

The computational strategy proposed in Griffiths *et al.*^[4]^ is the only attempt we are aware of that is aimed at estimating the rate and mitigating the effects of index hopping in droplet-based scRNA-seq data. Nonetheless, the strategy the authors propose, similar to other computational approaches we mentioned in the previous paragraph, does not in fact estimate the index hopping rate of individual reads, but rather computes a proxy measure, *the swapped fraction* in the case of Griffiths *et al.*^[4]^, which they define as the fraction of molecules (i.e. those with an identical UMI, cell barcode, gene label) that are observed in more than one sample. As we show in this paper, the probability at which an individual sequencing read swaps the sample index is not the same as the probability that one out of all the PCR-duplicated reads from the same molecule swaps the sample index. As we show, even when the sample index hopping probability at the level of individual read is very low, the fraction of *phantom molecules*, molecules that we observe to have swapped sample indices, can vary greatly, even as far as comprising the vast majority of molecules in a given sample. As for correcting for the effects of index hopping, the proposed molecular exclusion heuristic the authors propose suffers from high positive and negative rates, resulting in unnecessarily discarding molecules that otherwise should have been retained and conversely, in retaining molecules that should have been discarded. In particular, by overlooking the possibility that a given fraction of observations consists entirely of phantom molecules, the molecule exclusion strategy they propose can potentially retain a high proportion of false positives molecules. Furthermore, by assigning the true sample of origin as the sample with the largest read fraction while discarding all molecules that have a read fraction below a given threshold (default being ≤ 0.8), their strategy would further result in high false positive since neither the information implicit in the absolute read count distribution nor the extremely variable distribution of molecular proportions (i.e. *library complexity*) across samples and *PCR duplication levels* are considered. For example, even if an observation such as ***y*** = (1, 4, 0, 0) has the highest read fraction at *s* = 2, it would still be an unlikely event if for instance we have 70 times more molecules in Sample 1 than in Sample 2, that is, *π*_5_ = (0.7, 0.01, 0.19, 0.1).

##### Contrasts with Proposed Approach

The probabilistic model we propose in this paper aims at capturing the distributional patterns of sequencing read counts in order to both estimate the fundamental quantity of interest *the sample barcode index hopping rate* at the level of individual reads and quantify its manifestation as a data contamination artifact as measured by *the fraction of phantom molecules*. As we show (see Equation 8), the fraction of phantom molecules is determined not only by the sample index hopping rate, but also by the number of multiplexed samples, as well as by their library size, coverage, and molecular complexity profiles (i.e. the distribution of molecular proportions conditional on the *PCR duplication levels*). As such, this complex relationship, which we have attempted to capture in our proposed model, can potentially provide an explanation for the large variance in the index hopping rate estimates (0.2% - 8%) that have been reported in the literature^[3,4,7,11]^. Furthermore, we use the probabilistic modeling approach to discard molecules not based on their observed read fraction, but rather on a corresponding posterior probability that is a function of not only the distribution of the read counts, but also the model’s estimated sample index hopping rate, the library sizes and the expression profiles of the multiplexed samples, as the following paragraph illustrates.

##### Illustrative Examples from Empirical Data

Consider the following scenarios encountered in the data. For example, in the HiSeq 4000 dataset, *q* (the maximum posterior probability of the true sample of origin) is higher for outcome *y*^(*opt*)^ = (0, 0, 0, 1, 1, 0, 0, 1) than it is for *y*^(*below*)^ = (0, 1, 2, 0, 0, 0, 1, 0) even though in the latter, the inferred true sample of origin *s* = 3 (Sample B1) has two reads whereas the three molecules in the former has one read each, including the inferred true sample of origin *s* = 5. This might seem counter-intuitive at first sight, but once we consider the proportion of molecules, it becomes apparent why it makes sense to retain the molecule corresponding to *s* = 5 but discard the one corresponding to *s* = 3. For this observation, we have 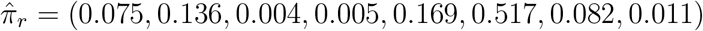 for *r* = 3 for *r* = 4. Given that the lowest proportions are for *s* = 3, 4, 8 in that order, and highest for *s* = 5, it does make sense indeed to expect that the two reads *y*^(*opt*)^ originated from *s* = 5. In contrast, for *y*^(*below*)^, it is not really clear that the hopped reads originated from *s* = 3 given the sample has the lowest proportion of molecules. For the HiSeq 2500 data, we have 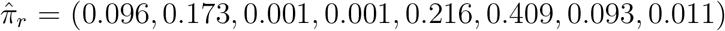 for *r* = 3, which explains why we end up discarding the molecule *s* = 8 in *y*^(*below*)^ but not in *y*^(*opt*)^ since there are half the number of molecules in *s* = 5 than in *s* = 6, making it more likely that the read indeed hopped from *s* = 8. We have a similar situation for the NovaSeq 6000 datasets where the sample with the lowest proportion of molecules (i.e. 0.005) corresponds to *s* = 15 (i.e. Sample *P*7_8), making it more likely that the two hopped reads in *y*^(*opt*)^ of L1, the one hopped read in in *y*^(*below*)^ of L1, and the one hopped read in *y*^(*opt*)^ of L2 actually originated from *s* = 9. This also makes *y*^(*below*)^ of L2 more likely to be a fugue non-chimeric observation (see definitions, Methods Section 3.6), such that the one read in *s* = 15 would in reality have had hopped from a sample with a larger number of molecules.

## 3 Methods: Model Formulation

### Scope

Although much of what we propose here can be ported, modified, and applied to other protocols, we limit the scope of the proposed approach to droplet-based scRNA-seq data, with libraries prepared using the 10x Genomics Single Cell Protocol and that are subsequently multiplexed for sequencing on Illumina machines.

### 3.1 Sequencing Reads Annotation

Consider a droplet-based single-cell sequencing experiment where a total of *S* (ranging from 2 to 96) libraries are pooled together and multiplexed on the same lane of a patterned flow cell. In a single sequencing run, millions of short sequencing reads are generated (from 350M reads on a single lane of HiSeq 4000 to 2.5B reads on a single lane of a NovaSeq 6000 S4 flowcell), each of which is annotated with barcodes representing the sample, cell, and molecule from which the read originated. If the reads are aligned to the genome, a read would also be annotated with the genomic location where it mapped to. That is, after sample demultiplexing and transcriptome alignment, each read becomes associated with a *cell-umi-gene-sample* label. More precisely, Each mapped read in a 10x Genomics Single Cell 3’ v2 Gene Expression Library can be annotated by four labels: (1) *A sample barcode*, (2) *cell-barcode index*, (3) *Unique Molecular Identifier* (UMI) (4) *gene ID*.

1. *A sample barcode*. In the case of a 10x Genomics single-indexed library, there are four indices for each sample. In what follows, we collapse all four indices into a single sample index for computational, mathematical, and practical considerations. However, as we show in Section 1.5, the model we propose allows us to recover the sample index hopping rate at the level of the barcodes.
2. A 16bp *cell-barcode index* randomly selected out of a set containing 737,280 possible combinations.
3. A 10bp *Unique Molecular Identifier* (UMI) index for which there are a total of 4^10^ = 1, 048, 576 combinations. A *UMI collision* occurs when two or more UMIs possess the same sequence.
4. A mapped *gene ID* as provided by a transcriptome. For example, Ensemble 95^[14]^ has annotations for 19,768 and 21,823 non-conjoined coding and noncoding genes, respectively.

#### UMI Collisions

Given that the UMI index in v2 chemistry is 10bp long, the number of possible UMIs is one order of magnitude larger than the number of UMIs typically observed in any one cell. Accordingly, a UMI collision in a single cell, when it does occur, would result in two reads from different molecules to be considered as originating from the same cDNA fragment (i.e. PCR duplicates), thus reducing the number of real molecules that can be observed in any one cell. To further reduce the chance of collisions, gene ID labels can be considered in order to limit the space of collisions to those UMIs that are mapped to the same region of the genome. Although these genomic regions, each consisting of the union of exons belonging to a given gene, vary in length, the localization of possible collisions to a small region of the genome greatly reduces the probability of UMI collisions.

#### RNA-containing cells

In scRNA-seq data, a cell is identified by a unique cell barcode. However, only a fraction (1k-10k) out of the +100K observed cell barcodes correspond to actual cells. In each of these cells, we typically observe anywhere from a minimum of a 1000 (a threshold usually specified manually) up to approximately 100,000 unique molecules, or more depending on the average read coverage per cell.

#### Discarding non-exon mapped reads

We note here that by discarding unmapped reads and reads that mapped to genomic regions other than exon bodies, we are further making the implicit assumptions that the large portion of data that we end up retaining contains enough information to determine the index hopping rate and that the data we discard is not characterized by a different mechanism underlying the sample barcode index hopping phenomenon.

### 3.2 Modeling Sequencing Reads

We start by observing that after cDNA molecules of a given transcript are amplified, they are subsequently fragmented into 300-400 bp fragments, each containing both a cell barcode and a UMI. During the sample index PCR amplification step, an Illumina Read 2 adapter is ligated to a fragment *m* before it gets amplified, resulting in *n*_*m*_ sequencing reads having the same *cell-umi-gene-sample* label. If none of the *n*_*m*_ sequencing reads are misassigned, then each *n*’th sequencing read’s observed sample index *d*_*mn*_ would correspond to the true sample index *s*_*mn*_. When index hopping occurs, the read misassignment process can be modeled as a mixture of *S* categorical distributions where each observed molecule’s read sample index belongs to a given sample. See Table 1 for an example toy data table in which each row corresponds to a single sequencing read with label *m*.

**Table 1:** Toy dataset I. Table of three multiplexed samples (i.e. *S* = 3) in which each observation is a sequencing read with label *m* (i.e. a unique combination of *c*, *u*, and *g*, which denote the cell-barcode, UMI, and gene, respectively), a true source sample *s*, and an observed sample *d*_*i*_, *i* = 1, 2, 3. In a typical experiment, a single lane can generate anywhere from 350 million to 2.5 billion sequencing reads.

| *label* | *l* | *s* | *d*<sub>1</sub> | *d*<sub>2</sub> | *d*<sub>3</sub> |
| --- | --- | --- | --- | --- | --- |
| $c_1u_2g_1$ | 1 | 1 | 0 | 1 | 0 |
| $c_1u_1g_1$ | 2 | 2 | 1 | 0 | 0 |
| $c_1u_3g_2$ | 3 | 2 | 1 | 0 | 0 |
| $c_1u_3g_2$ | 3 | 2 | 1 | 0 | 0 |
| $c_1u_3g_2$ | 3 | 2 | 0 | 1 | 0 |
| $c_2u_1g_1$ | 4 | 1 | 0 | 1 | 0 |
| $c_2u_1g_1$ | 4 | 1 | 0 | 0 | 1 |
| $c_2u_1g_1$ | 4 | 1 | 0 | 0 | 1 |
| ⋮ | ⋮ | ⋮ | ⋮ | ⋮ | ⋮ |

We can formulate the mixture model formally as a two-stage hierarchical sampling process, where we first draw a molecule with an unobserved true sample index *s*_*mn*_ that gets assigned to read *n* with label *m* from a categorical distribution governed by a label-specific probability parameter vector ***ϕ***_*m*_. Next we draw the observed sample index from another categorical distribution with a probability parameter vector ***p***_*smn*_ whose value is conditional on the value of *s*_*mn*_. In what follows, *s*_*mn*_ denotes the index of the element in the vector ***s***_***mn***_ for which we observe a one (i.e [***s***_***mn***_ = 1]).

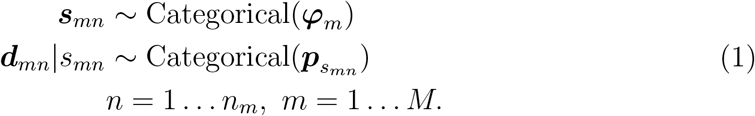

Where ***ϕ***_*m*_ ∈ [0, 1]^*S*^, ||***ϕ***_*m*_||_1_ = 1, 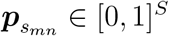, 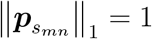, *d*_*mn*_ ∈ {1, …, *S*}. *s*_*mn*_ ∈ {1, …, *S*}, *S* ∈ ℕ_+_.

#### The sample hopping probability matrix

The probability parameter vectors of all the *S* distributions can be stacked together in ***P*** as such.

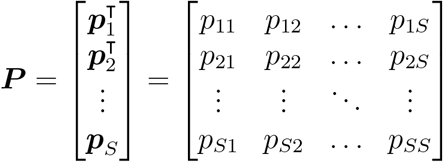

When ***P*** = **I**, then the probability that a read keeps its sample index of origin is one, and consequently, reads do not hop over to other samples. That is, *d*_*mn*_ = *s*_*mn*_ for all *m* and *n*. Here *p*_*ij*_ denotes the probability that a read from sample *i* hops to sample *j*. The number of parameters in ***P*** equals *S* ×(*S* − 1). Furthermore, in Model 1 there are a total of *M* parameter vectors ***ϕ***_*m*_ each taking values on the probability simplex for a total of *M* × (*S −* 1) parameters. We can reduce the number of parameters from (*M* + *S*) *×* (*S −* 1) to 1 by making the following two assumptions.

### 3.3 Two Simplifying Assumptions

The *n*_*m*_ reads associated with label *m* can come from different source samples. This could happen for example when reads from two samples are assigned the same label simply by chance. However, such an outcome is extremely unlikely and by ruling it out we can greatly simplify the modeling framework if we constrain each of all the *M* parameter vectors ***ϕ***_*m*_ to be the one vector (i.e. **1**). More formally, we formulate the assumption as follows.

#### Assumption 1 (Discrete Simplex Constraint Assumption)

Let *max*(***ϕ***_*m*_) = 1 for all *l* such that 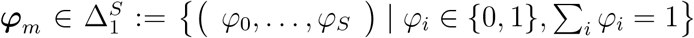. That is, we restrict ***ϕ***_*m*_ to belong to the discrete *S*-simplex, a set with cardinality *S* whose elements are vectors each consisting of a single 1 and zeros elsewhere. We basically assume that all the reads with a given *cell-umi-gene* (CUG) label (i.e. PCR read duplicates of a unique cDNA molecule) can be indexed with the same sample barcode only.

As a result of Assumption 1, the sample index *s*_*mn*_ becomes a constant random variable drawn from one of *S* degenerate categorical distributions such that all the *n*_*m*_ reads with label *m* are assigned the same sample index. That is, for each ***ϕ***_*m*_, all the 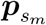 vectors become the same for all the *n*_*m*_ reads (i.e. 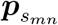). Assumption 1 entails the following two corollaries.

#### Corollary 1 (Label Collision Across Samples)

A given *cell-umi-gene* label combination has a zero probability of co-occurring with more than one sample barcode index. That is, reads annotated with the same label combination can only belong to one sample.

#### Corollary 2 (Number of Molecules)

The number of total molecules across all *S* samples equals the number of unique *cell-umi-gene* label combinations *L*. That is, each CUG would represent one unique molecule and all sequencing reads with the same CUG would correspond to PCR amplification products of that original molecule. In other words, the number of molecules equals the number of rows in a merged data table of sample read counts, which has been fully joined by the combination of cell, umi, and gene key.

Furthermore, given that we have no reason to believe that reads in a particular sample are characterized by a different chemical properties, we can simplify the problem by making a second assumption.

#### Assumption 2 (One parameter to rule them all)

The probability of a read keeping its sample index is the same across samples (i.e. *p*) and the probability of a read switching the sample index is the same regardless of either its source or target sample (i.e. 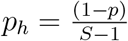).

Assumption 2 reduces the number of parameters in *P* from a total of *Q* = (*S* − 1) × *S* to just a single parameter.

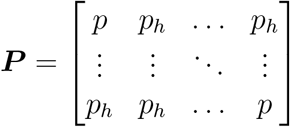

More importantly, Assumption 2 induces a symmetry on the probabilities we assign to the possible counts that lie within the same *m*–face of an (*S* − 1)–simplex denoted by 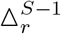, which in turn constitutes the *r*’th component of the Pascal’s Simplex denoted by *∧*^*S*^.

### 3.4 Modeling Sequencing Reads Counts

Assumption 1 allows us to sum over the *n*_*m*_ categorical random variables to obtain a mixture of multinomials model for data {***y***_*m*_}, where ***y***_*m*_ = (*y*_*m*1_,…, *y*_*ms*_,…, *y*_*mS*_) denotes the vector of read counts across *S* samples corresponding to an observed CUG label (i.e. molecule) *m* = {1, …, *M*} originating from an unobserved source sample 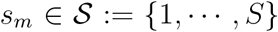. Here, the total number of CUGs (i.e. molecules) is *M* ∈ ℕ_+_; the total number of PCR duplicated reads associated with molecule *m* is by ||***y***_*m*_||_1_ = *n*_*m*_; and the total number of mapped reads across all samples in the dataset is *N* = ||***n*** = (*n*_1_,…, *n*_*M*_)||_1_ where ***n*** is the *M* -dimensional vector of read count sums.

We are interested in specifying a model of read counts conditional on a given unique PCR duplication level. So in what follows, we partition ***n*** by the unique values that *n*_*m*_ can take. That is, the number of PCR duplicated reads -whether hopped or not - a given unique molecule generates, more specifically, 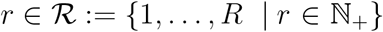, where 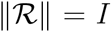 and *R* is the maximum value in 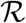. For each 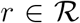, let 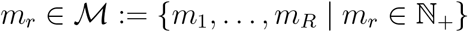 denote the corresponding number of times *r* is observed in ***n*** such that 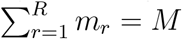. We denote the empirical distribution of *r* by 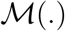. Accordingly, we can write the mixture of multinomials model conditional on the *PCR duplication level r* as follows.

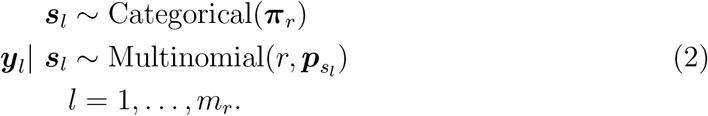

where *l* now indexes the *m*_*r*_ observations with *PCR duplication level r*; 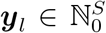, *S* ∈ ℕ_+_, ||***y***_*l*_||_1_ = *r*, *s*_*l*_ ∈ *s*, ***π***_*r*_ ∈ [0, 1]^*S*^, ||***π***_*r*_||_1_ = 1, ***p***_*s*_ ∈ [0, 1]^*S*^, ||***p***_*s*_||_1_ = 1; the vector 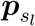 denotes the *s*_*l*_ row of the *S* × *S* sample hopping probability matrix *P*; and the probability vector ***π***_*r*_ represents the proportion of molecules across the *S* samples at *PCR duplication level r*.

The mixture model can be viewed more intuitively as a generative process in which we first sample a molecule ***s***_*l*_ from a library sample *s*_*l*_ according to the categorical model then we amplify the molecule by generating *r* PCR read duplicates according to the multinomial model. The number of molecules originating from a given sample with a given PCR read duplicates *r* is determined by the parameter vector *π*_*r*_ whereas the number of PCR duplicated reads that end up hopping to other samples is determined by the parameter vector 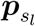).

### 3.5 The Molecular Proportions Complexity Profile

In a multiplexed experiment, several samples that vary in their library complexity are sequenced together, where we define *library complexity* as the expected number of unique molecules sampled with a finite number of sequencing reads generated in a given high-throughput sequencing run. These samples would differ in the total number of unique transcripts each one has due to a host of factors, ranging from the presence of varying amounts of RNA that characterize different cell types (e.g. neuronal cells have low RNA content) to accidental errors in library preparation that could cause many cells to break up and lose their endogenous mRNA. That is, even if total number of available sequencing reads were budgeted evenly over the multiplexed samples, the number of unique molecules detected in the sequencing run could vary widely across the samples. In order to assess and identify potential problems such as low library complexity across all the samples in a sequencing run, we propose that a more informative picture could be gained into the root cause of a sample’s, if we consider the molecule counts conditional on the *PCR duplication level r* or more specifically, the set of molecular proportions 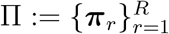, which we term the *molecular proportions complexity profile* (see Fig. 4).

We can obtain an estimate for *π*_*rs*_ from the observed proportion of read counts 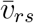 for sample *s* observations at *PCR duplication level r* by the following formula (for derivation see **Supplementary Notes 1.1**).

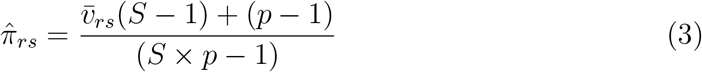

### 3.6 Definitions

Corollary 2 implies that since a distinct molecule is defined by a read (or multiple reads) with a unique *cell-umi-gene* label, any label collision across samples would result in a total number of *observed* molecules greater than *M*. Therefore, to avoid any potential naming confusions, we make the following clarifying definitions.

#### Definition 1 (Chimeric Observation)

A chimera refers to a hybrid creature composed of the parts of more than one animal. We use the analogy to refer to those read count *observations* for which 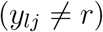 for all *j* = 1, …, *S*, or in other words, to those *cell-umi-gene* labels where we observe reads from more than one sample. We can further classify chimeras by the number of collisions. A k-chimera has reads in *k* sample categories (e.g. *y* = (4, 1, 0, 2, 0) is a 3-chimera when *S* = 5). In contrast, we refer to an observation with no collisions (i.e. a 1-chimera) as a non-chimera.

If there is no sample index misassignment whatsoever, then we expect that all the *M* observations to be non-chimeric. When the sample index hopping rate is large, we expect to see a correspondingly large proportion of chimeras, observations where there are collisions of labels across the samples.

#### Definition 2 (Phantom Molecule)

For a given observation with label *cell-umi-gene*, we term molecules observed in samples other than the true source sample as *phantom molecules*. We call them *phantom* since their existence is not real, due only to an artifact brought about by index hopping.

It is important to note that a sample index can hop even when we do not observe a label collision. Thus one or even all reads annotated with a given label may be actually hopping reads originating from the true source sample. In such a case, we are unable to determine the true source sample of these molecules. Nonetheless, we can estimate their expected observed proportions.

#### Definition 3 (Fugue Observation)

A disassociative fugue is a personality disorder characterized by unplanned travel, wandering, and even the establishment of a new identity. Similarly, for a given label, when all the reads hop over from an unobserved sample to establish a new identify in other samples, we term all such observations as *fugues* since we cannot know where they came from since we observe zero reads associated with the true sample of origin.

#### Definition 4 (Sample Index Hopping Rate)

Whereas the term 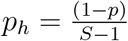 represents the probability that a sample index hops into a one particular target sample, the complement of *p*, namely 1 − *p*, refers to the quantity of interest, the Sample Index Hopping Rate, which we abbreviate as **SIHR**.

### 3.7 Toy Data Example

To illustrate the concepts and definitions in the preceding sections, consider the following toy example where we have three samples *S* = 3. The data would thus consist of three-dimensional discrete vector observations 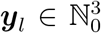 sampled from a multinomial distribution with total sum of counts *r*, where ||***y***_*l*_||_1_ = *r*, and a probability parameter.

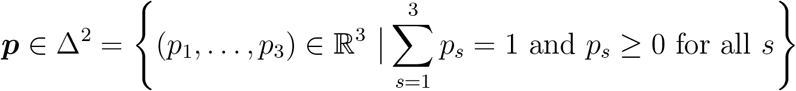

For concreteness, assume that the source sample *s* is known for each observation and consider what the data would look like when the sample index hopping rate (i.e. probability) is not zero (e.g. ***p***_1_ = [.9,. 05,. 05]) such that the probability concentration bleeds into outcomes with hopped reads (see Table 2). That is, we see that the same label we associate with a unique molecule co-occurs with more than one sample index. When there is no index hopping, for example, when **SIHR**= 0 (e.g. ***p***_2_ = [0, 1, 0]), we observe reads only in one of the three samples (i.e. *non-chimeric non-fugues*). In Table 2, we show observations corresponding to only 4 out of the 10 possible outcomes at read count level *r* = 3. However, when *r* is large, the set of possible outcomes increases exponentially. When *S* = 3, probabilities of observing these outcomes are given by the corresponding terms of the trinomial expansion of ***p***.

**Table 2:** Toy dataset II. Data table of read counts and corresponding deduplicated counts (i.e. molecules). The data table shows 4 out of the 10 possible outcomes at read count sum level *r* = 3 for multiplexed data with *S* = 3 samples, see Figure 3. All 4 observations are associated with a unique *cell-umi-gene* label *l* each and an unobservable true sample of origin *s* = 1. For each read count ***y***, a vector of deduplicated read counts or molecules ***x*** is given. The chimeric number *k* denotes the number of molecules observed in each outcome and *f* denotes the number of phantom molecules. Although the chimeric vs. non-chimeric state of an observation can be directly seen from the data, the fugue vs. non-fugue state of an observation cannot be directly inferred, since it depends on knowing the true origin of the reads (i.e. the latent variable *s*). When there is no index hopping, the only outcomes possible are those classified as non-chimeric non-fugues.

| | | | | | $\mathbf{y}$ (reads) | | | $\mathbf{x}$ (molecules) | | | category |
| --- | --- | --- | --- | --- | --- | --- | --- | --- | --- | --- | --- |
| $l$ | $r$ | $k$ | $f$ | $s$ | $y_1$ | $y_2$ | $y_3$ | $x_1$ | $x_2$ | $x_3$ | |
| 1 | 3 | 1 | 0 | 1 | 3 | 0 | 0 | 1 | 0 | 0 | non-chimeric non-fugue |
| 2 | 3 | 1 | 1 | 1 | 0 | 3 | 0 | 0 | 1 | 0 | non-chimeric fugue |
| 3 | 3 | 2 | 1 | 1 | 1 | 2 | 0 | 1 | 1 | 0 | chimeric non-fugue |
| 4 | 3 | 2 | 2 | 1 | 0 | 1 | 2 | 0 | 1 | 1 | chimeric fugue |
| ⋮ | ⋮ | ⋮ | ⋮ | ⋮ | ⋮ | ⋮ | ⋮ | ⋮ | ⋮ | ⋮ | ⋮ |

To better visualize the distribution of possible outcomes, consider the case at the read count level *r* = 3 when reads hop from one sample only, that is, when the true sample of origin is *s* = 1 (See Fig. 3). When *r* = 3 the number of coefficients and thus corresponding outcomes (i.e. elementary events) are given by the (*r* + 1)‘th *triangular number* 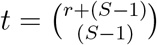, which here equals 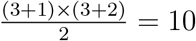. That is, we group the 10 outcomes, which make up the sample space 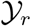 into a set of three categories: (1) Three non-chimeric outcomes, each lying on one of the three 0*−*faces (vertices); (2) Six 2*−*chimeric outcomes, two lying on each of the three 1-faces (edges); (3) One 3 − chimeric outcome lying in the middle of a single 2 − face (facet). In general, the number of *m*-faces of an *r*-simplex is 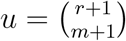. The unit probability measure is split over *r* outcomes instead of *t* outcomes. Out of the *u m*−faces, 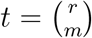 correspond to non-fugue outcomes whereas *f* = (*u* − *t*) are fugues (i.e. those observations where all the reads hop from their source sample). That is, if in Fig. 3, the true sample of origin is *s* = 1, then a 1-read hop would correspond to two possible outcomes: two 2-chimeric observations each giving rise to one true and one phantom molecule; A 2-read hop would correspond to three possible outcomes: two 2-chimeric observations each giving rise to one true and one phantom molecule and a single 3-chimeric observation giving rise to one true and two phantom molecules; A 3-read hop would would correspond to four possible outcomes: two 2-chimeric fugue observations each giving rise to two phantom molecules and a two non-chimeric fugue observations, each giving rise to one phantom molecule. Note however that when we consider hopping from all *S* samples, the number of possible outcomes *e* increases by a factor of *S*, so when *r* = 3, we have *e* = 30 possible outcomes.

**Figure 3:**
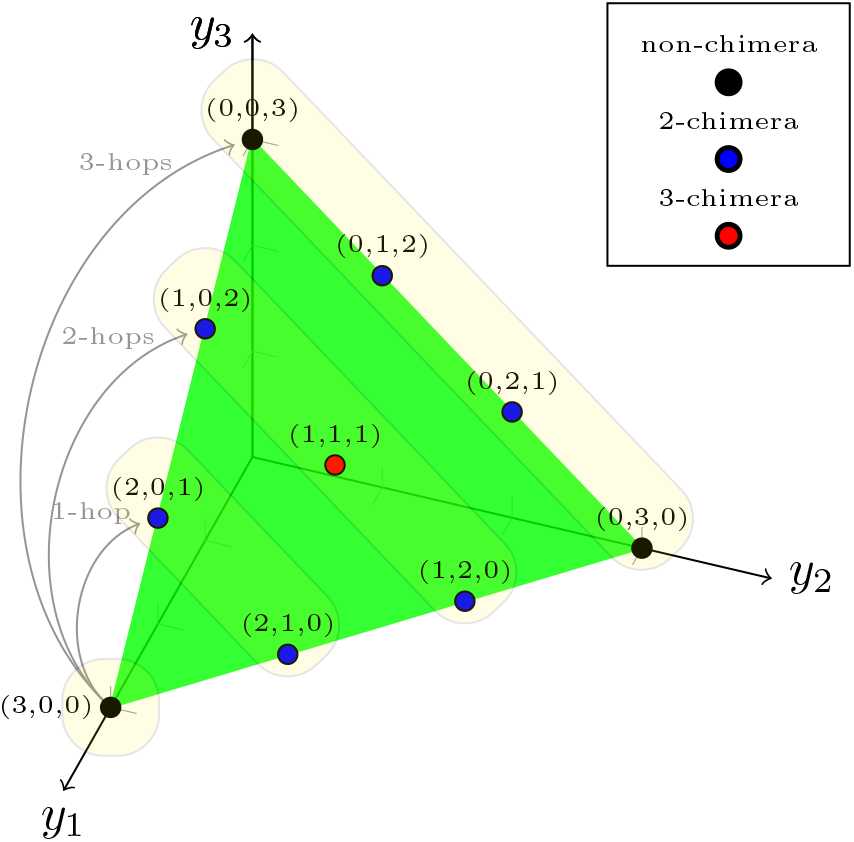
The 3rd component of Pascal’s 3-Simplex denoted by 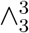. All 10 possible outcomes (at read count sum r=3) for three samples (S=3) are arranged on the simplex. The sample of origin is *s* = 1 as shown by the arrows. Outcomes can be classified according to the number of observed molecules (i.e. k-chimera) or the number of hopped reads. The four outcomes corresponding to all 3 reads hopping over are called *fugues*. The two fugue outcomes in the vertices are termed *non-chimeric fugues* whereas the other two on the edge are termed *chimeric fugues*

The number of observations in data 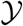 is typically high (in the hundreds of millions). Each observed vector of read counts can be categorized as a combination of *chimeric/non-chimeric* and *fugue/non-fugue*. For each k-chimeric observation, we potentially have from one and up to *S* − 1 *phantom molecules*. For example, the observation ***y***_4_ has two *phantom molecule* and zero real molecules. If there is indeed no index hopping, then the number of *phantom molecules* would be zero whereas the number of real molecules would equal to *L*, the number of unique label combinations. Although the probability that all reads get misassigned to the same sample is negligibly low, the effect can be significant for non-chimeric labels for which there is only one or two reads (a group that makes up a large fraction of observations).

## 4 Methods: Estimation of the Sample Index Hopping Rate

Given the large number of observations (i.e. number of CUGs *M*) that are commonly encountered in practice, we proceed to simplify the problem analytically to make computation tractable. In what follows, we reduce the mixture model to a single parameter model for the distribution of non-chimeric observations, we then derive the distribution of the sum of non-chimeric observations and show, after making a third simplifying assumption, that the distribution’s mean function corresponds to the solution of a differential equation governing a negative growth process. Then, we show how the index hopping rate can be estimated by formulating the problem as generalized linear regression model for binomial counts with a log link function that corresponds to the solution of the differential equation.

### 4.1 Modeling the Distribution of k-chimeras

The distribution of k-chimeras is essentially the distribution of the total number of non-zero counts of a multinomial random variable. With regards to single cell data, determining whether a sample count is nonzero is equivalent to a deduplication process of obtaining molecule counts from read counts. Statistically, deduplication can be formulated as a thresholded latent variable model where each element of a potentially unobservable random vector ***y***_*l*_ is thresholded into an observable Bernoulli random variable *z*_*ls*_ with an indicator function 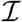 that detects whether a molecule is observed. That is,

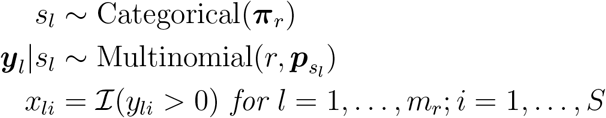

where 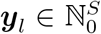, ***p*** ∈ [0, 1]^*S*^, ||***p***||_1_ = 1, and *S* ∈ ℕ. The marginal distribution of each element of the multinomial observation ***y***_*l*_ is binomial.

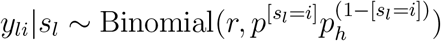

where the Iverson bracket notation is used. As we show in **Supplementary Notes 1.2**, the Bernoulli random variables (i.e. the elements of ***x***_*r*_) can be treated as independent (for *r* > *S*) but not identically distributed, and their sum, which indicates the category of the observation (i.e. *k*-chimera), can be given by the Poisson Binomial distribution.

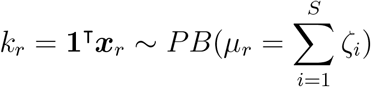

### 4.2 Modeling the Distribution of Non-chimeras

For the case of *k* = 1, or non-chimeric observations, we can derive a closed form of the distribution by noting that a *non-chimera* is a count observation ***y***_*l*_ for which (*y*_*li*_ = *r*) for any *i* ∈ {1, …, *S*}. We denote the event of observing a *non-chimera* by a Bernoulli random variable 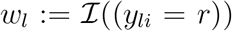 with mean parameter given by 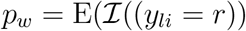. As a result, the distribution of observing a *non-chimera* is given by

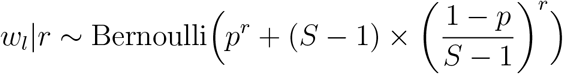

where 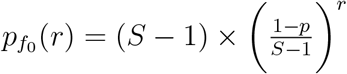 is the probability of observing a non-chimeric fugue observation with *r* reads and *p*^*r*^ is the probability of observing a non-chimeric non-fugue observation with *r* reads.

### 4.3 Modeling the Distribution of Sum of non-Chimeras

Given that *p*_*w*_ is the same for all observations with the same *PCR duplication level*, we can sum all the *m*_*r*_ Bernoulli random variables in *PCR duplication level r* to obtain a Binomial distribution over the number of *non-chimeras z*_*r*_ conditional on *r* **Supplementary Notes 1.3**. That is, for a given *r* we have

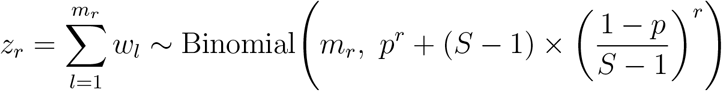

### 4.4 Estimating the Sample Index Hopping Rate

The joint sampling distribution of the chimeras at all *PCR duplication level* values, concatenated as a vector ***z***, can be decomposed as follows.

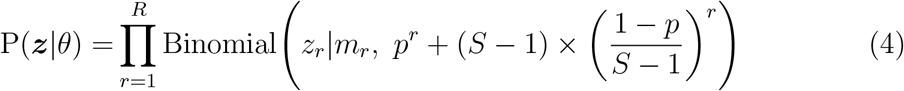

#### Assumption 3

(The Negligible Contribution of the Probability of Observing non-Chimeric Fugues). Given that the *the sample index hopping rate* we observe in experimental data tends to be very small, *SIHR* < 0.05, the contribution of the second term (i.e. the probability of non-Chimeric fugues) in the parameter of Model 4 is discernible only when *r* ≤ 2.

With the aid of Assumption 3, Model 4 can be simplified and the relationship between the number of non-chimeras *z*_*r*_ and the sample index hopping rate (1 − *p*) at a given *PCR duplication level r* and with an *m*_*r*_ number of observations can be formulated as a *generalized linear regression* model with a *log* link function as follows (see **Supplementary Notes 1.4** for details).

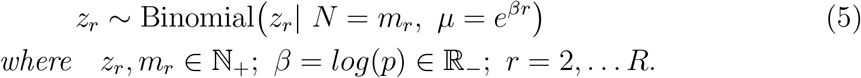

An estimate of the sample index hopping rate can be obtained from the regression coefficient.

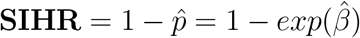

From which we can obtain the sample barcode index hopping rate (**Supplementary Notes 1.5**).

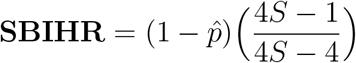

## 5 Methods: Reassigning Reads and Purging Phantom Molecules

### 5.1 Inferring the True Sample of Origin

In Model 2, we denote the true sample of origin of a sample index by the latent variable ***s_l_***, whose posterior probability distribution can be derived via Bayes theorem. The posterior distribution allows us to quantify our residual uncertainty about the true sample of origin given the observed read count and the set of parameters *p* and the molecular proportions 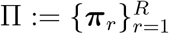 that govern the mixture model.

#### 5.1.1 The Posterior Distribution of the True Sample of Origin

The posterior distribution for each element is given by.

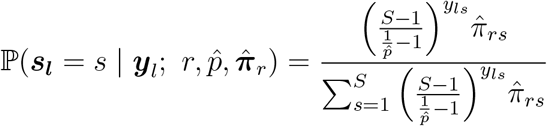

where the plug-in estimates 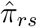 is the proportion of molecules in sample *s* at PCR duplication level *r* as computed by Equation 3 and 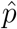 is the complement of the sample index hopping rate as estimated by Model 5. For derivation, see **Supplementary Notes 1.6**. We label the sample with the maximum posterior probability as the true sample of origin.

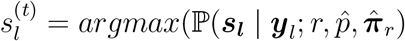

The corresponding posterior probability of 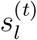 is simply the maximum over the posterior probability vector.

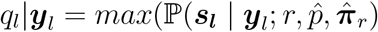

When 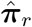 is uniform, the maximum *q*_*l*_|***y***_*l*_ can have duplicated values. However, such an outcome is extremely unlikely given the high variability of the molecular proportions that characterize empirical data and therefore will not be considered.

### 5.2 Purging Phantom Molecules

For each of the *L* observations, we label the molecule corresponding to 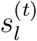 as a real molecule and all the others (with nonzero reads) as phantom molecules. Such a procedure achieves the minimum possible number of false negatives at the expense of false positives. However, it could be well the case that sacrificing a potential real molecule is worth more than retaining a phantom molecule in the dataset. In scientific applications, most often, a weak signal is preferable to an artifactual signal. That said, to achieve the minimum possible number of false positives, it might be the case that the entire data would need to be discarded. An alternative approach attempts to find the optimal trade-off that minimizes both the false positive and false negative rates.

To optimally minimize the false positive rate, we would need to find the optimal cutoff value *q*^∗^ below which, observations with 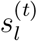 that we labeled as real molecules are now relabeled as phantom – in other words, we discard them along with previously labeled phantom molecules. In order to determine *q*^∗^, we would need to work with the marginal posterior cumulative distribution function of *q*, which does not have a closed-form expression but which can be expressed as such.

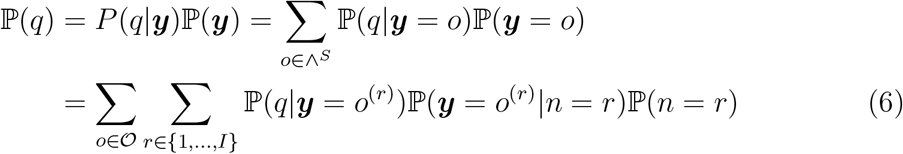

Here, 𝒪 consists of all the outcomes ***y*** that correspond to the coefficients in *r*’th component of the Pascal’s S-Simplex 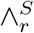. Whereas ℙ(***y*** = *o*^(*r*)^ *n*) is given by the multinomial distribution can be computed numerically for the first dozen values of *PCR duplication level r*, the marginal distribution of *r*, P(*n*), can be approximated by the observed proportion of *PCR duplication levels* which we denoted by *M* (.) in Model 2. However, since *r* can have large values in empirical data, we can work with the empirical distribution instead and approximate the marginal distribution of *P* (***y***) by the observed relative frequency distribution of outcomes (e.g. the proportion of ***y*** = (1, 2, 0, 0) observations in the data).

### 5.3 Determining the Optimal False Positive Rate

The approach outlined in the previous section can be thought of as a classification task in which we attempt to predict whether a given molecule is a phantom molecule, which we consequently discard, or a real molecule, in which case we retain. We would like to minimize both the error that a molecule we deem real is in fact a phantom (i.e. false positive) and the error that a molecule we deem phantom is in fact a real molecule (i.e. false negative). In a dataset consisting of *L* observations and *M* molecules, the maximum number of real molecules is *L*. In what follows, it helps to work with proportions relative to the number of observations *L*. Accordingly, we define 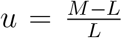 as the *molecule inflation factor*, a lower limit on the fraction of molecules we can predict to be phantom (with respect to *L*). Although the initial total number of molecules in the entire data before index hopping is *L*, the upper limit of the number of real molecules we are able to predict as real is slightly less than *L* due to the existence of fugue observations in the dataset, each one of which would lead to one extra true phantom molecule to be unaccounted for. We term these molecules as *fugue phantoms*. *L* therefore would equal the sum of real molecules and fugue phantoms. Accordingly, if *g* is the proportion of fugue observations in the dataset, which we can obtain as shown in **Supplementary Notes 1.8**, then there would be a total of (*u* + *g*) true phantom molecules and a total of (1 − *g*) true real molecules in the data. For example, for a dataset with *L* = 100 observation, a total number of molecules *M* = 150, and an estimated proportion of fugues *g* = 0.05, the *molecule inflation factor u* would equal 0.5 so that the total number of true phantom molecules is given by (0.5 + 0.05) × *L* = 55 and the total number of real molecules is given by (1 − 0.05) × *L* = 95.

Given that the number of phantom molecules in a given dataset is given by *L ×* (*u* + *g*), there are three courses of action we can choose to take.

1. *No Purging*: We keep the data as it is. By doing so, we basically label all the phantom molecules as real, which would correspond to *L* × (*u* + *g*) false positives (i.e. FPR=1). That is, we are in effect incorrectly classifying all true phantom molecules as real.
2. *No Discarding*: We purge the phantom molecules by reassigning the reads to the sample with largest posterior probability. By doing so, we can drastically decrease the number of false positives while incurring relatively small number of false negatives.
3. *Discarding Below Cutoff*: We can decrease the false positives further at the cost of a slight marginal increase in false negatives by choosing a cutoff *q*\* below which predicted real molecules are classified as phantom instead. That is, in our classification task, we label the molecule with the maximum posterior probability (*q*) above or equal to a given threshold *q*\* as a real molecule while the remaining molecules are all labelled as phantoms. Furthermore, all the molecules in observations whose corresponding *q* falls below the cutoff are also labeled as predicted phantoms even though a proportion of them are real molecules which we cannot confidently classify (i.e. false negatives). Effectively, all molecules we label as phantom get eliminated from the dataset.

In what follows, it would be easier to work with the complement of *q*, which we denote *qr* = 1 − *q*, which is the probability of a molecule originating from a sample other than the one with maximum posterior probability. For a given selected threshold value *qr*^∗^, we let *o*^∗^ = *F*_*qr*_(*qr*^∗^):= Pr(*qr* ≤ *qr*^∗^) denote the proportion of observations whose corresponding predicted real molecules we retain. Here, *F*_*qr*_ is the empirical CDF of *qr*. Consequently, the fraction of predicted real molecules is *o*^∗^ and the fraction of predicted phantom molecules is 1 + *u* − *o*^∗^, since the total fraction of molecules must sum up to *u* + 1 (see Table 3). Out of the (*o*^∗^) predicted real molecules, the proportion of false positives is given by the expectation of the probability of false assignment over the subset of the data *o*^∗^.

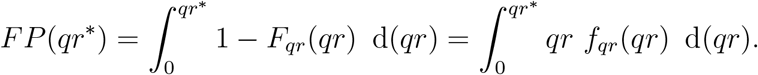

**Table 3:**
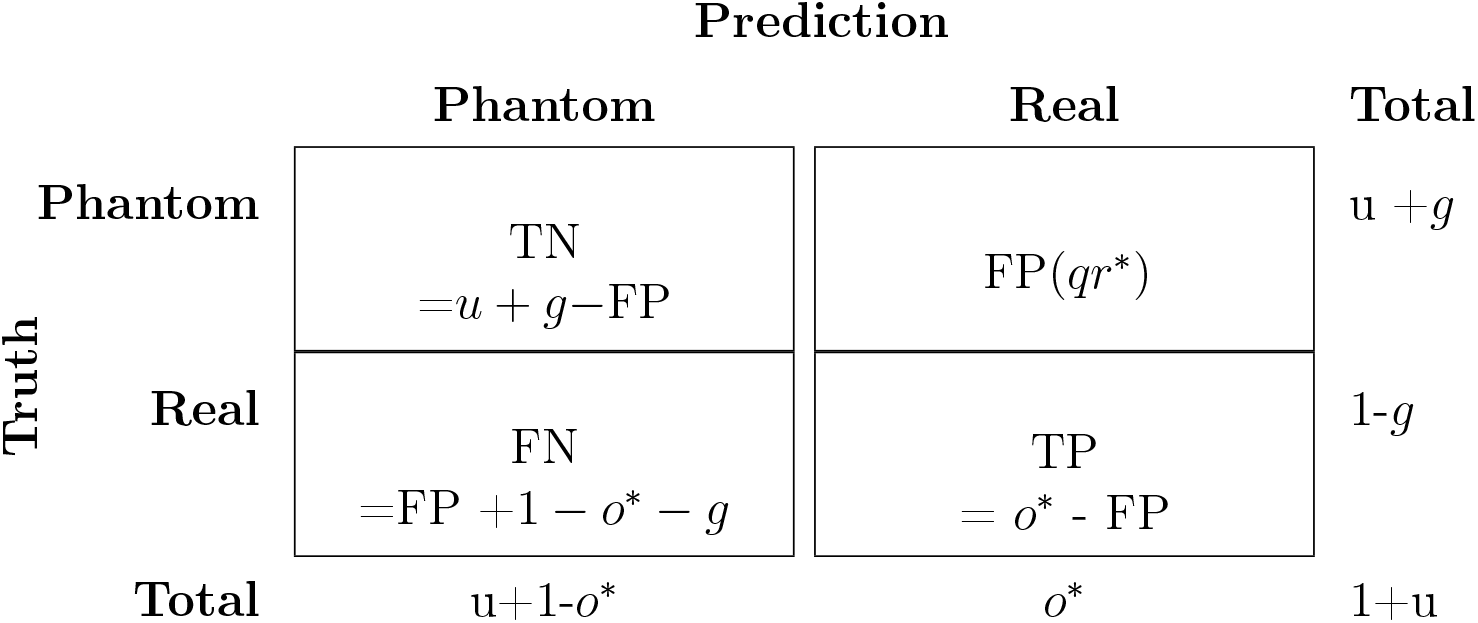
Confusion Matrix. Possible outcomes for classifying molecules as real or as phantom. Given a data table of read counts with *L* observations, the total number of molecules would be (1 + *u*) × *L* where *u* is the *molecule inflation factor*. The variable *o*^∗^ denotes the proportion of observations that fall above threshold *q*^∗^ and such that *o*^∗^ × *L* would be the number of molecules we label as real. The constant *g* is the proportion of fugue observations estimated for that given dataset.

|  |  | Prediction |  |  |
| --- | --- | --- | --- | --- |
|  |  | Phantom | Real | Total |
| Truth | Phantom | TN<br>$= u + g - \text{FP}$ | $\text{FP}(qr^*)$ | $u + g$ |
| | Real | FN<br>$= \text{FP} + 1 - o^* - g$ | TP<br>$= o^* - \text{FP}$ | $1 - g$ |
| Total | | $u + 1 - o^*$ | $o^*$ | $1 + u$ |

Note that the empirical cumulative distribution function of $qr$ is discrete (i.e. its ECDF increases by jump discontinuities only) even though it takes real values in $\mathbb{R}^{[0,1]}$ . Accordingly, the threshold $qr^*$ should be set such that very few potential real molecules are discarded for any marginal decrease in false positives.

Note that the empirical cumulative distribution function of *qr* is discrete (i.e. its ECDF increases by jump discontinuities only) even though it takes real values in ℝ^[0,1]^. Accordingly, the threshold *qr*^∗^ should be set such that very few potential real molecules are discarded for any marginal decrease in false positives.

#### TORC approach

The optimal value for the threshold *qr*^∗^ is the one that simultaneously minimizes both the number of FP counts and FN counts, but note that discarding lower quality data would lead to a small FP but also a large FN. Although we can use *Youden’s***J** statistic = 1-(FPR +FNR) to find the optimal cut-off value *qr*∗ that maximizes **J**, where 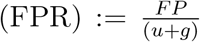 and 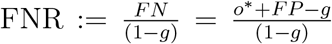, the measure is not appropriate for our application (see **Supplementary Notes 1.7**). An alternative approach for minimizing both the number of FP counts and FN counts considers the ratio of marginal increase in FNs over the marginal decrease in FNs as a trade-off to be specified by the researcher, the Trade-Off Ratio Cutoff *TORC* parameter, which represents the maximum number of real molecules one is willing to incorrectly discard in order to correctly purge one phantom molecule. That is, we make our reference coordinates (the point of origin) to be the number of false positive and false negative molecules we obtain if we reassign reads without purging any predicted real molecules. As we discard an increasing number of predicted real molecules, the marginal false positive (i.e. predicted real molecules that are actually phantom) decreases while the marginal false negative (i.e. real molecules that we effectively predict as phantom by discarding) increases. For a given dataset, the cutoff *TORC\** that gets effectively chosen would correspond to the largest observed *TOR* value not exceeding the preset *TORC* value. All molecules with corresponding *TOR* values strictly less than *TORC\** -not *TORC* -are discarded. For example, if we have *tor* = (0.1, 0.5, 2.9, 4.1, …) and *TORC=3*, then *TORC\** =2.9 and predicted real molecules corresponding to *tor* = 0.1 and *tor* = 0.5 are discarded.

## 6 Data Analysis

### 6.1 Data Preprocessing

#### Empirical data

We applied the proposed model on three publicly available 10X Genomics scRNA-seq datasets: (1) a control non-multiplexed dataset in which each sample was sequenced on a separate lane of HiSeq 2500; (2) a multiplexed dataset sequenced on HiSeq 4000; and (3) a multiplexed dataset sequenced on NovaSeq 6000. The HiSeq 2500 and HiSeq 4000 datasets consist of 8 libraries of mouse epithelial cells, which have also been used previously by Griffiths *et al.*^[4]^ for analysis of sample index hopping. The two datasets were downloaded from the authors’ host server using the *get_data.sh* script available on our paper’s GitHub repo (for a more detailed description of the data, please refer to the original publication,^[1]^). The third dataset was obtained from the Tabula Muris project’s repository. It consisted of 16 libraries (i.e. the P7 libraries) of mouse tissue samples, which were pooled and multiplexed on two lanes of an S2 flowcell in a single NovaSeq 6000 sequencing run^[2]^. BAM files for the 16 samples were downloaded from the NIH’s SRA data repository (SRA accession number SRP131661) and converted back to FASTQ files using the 10X Genomics *bamtofastq* utility. Cell Ranger 3.0.0 was run with the default options on each set of samples multiplexed on the same lane.

#### Validation Data

Two 10X Genomics scRNA-seq sample libraries were sequenced in two conditions. In the first condition, the samples were multiplexed on the same lane. In the second condition, two sample libraries were sequenced on two separate lanes of HiSeq 4000 (this non-multiplexed condition provides a ground truth for the true sample of origin of each CUG). Cell Ranger 3.0.0 was run with the default options on each of the four samples (two sample multiplexed on the same lane and the same two samples non multiplexed. A joined read counts table was created and gene IDs anonymized. The joined table had 16,547,728 unique CUGs, out of which 9,252,147 CUGs were present in both the multiplexed and non-mulitplexed samples. These were retained and a column containing goundtruth labels was added to the resulting inner joined datatable. Rows corresponding to colliding CUGs were filtered out and the table saved to file (see the reproducible R markdown notebook *validation_hiseq4000_1.nb.html* for details). Both read counts tables was deposited on Zenodo (https://doi.org/10.5281/zenodo.3267922) and linked on the GitHub project’s website.

### 6.2 Molecular Proportions Complexity Profiles

Figure 4 depicts the molecular proportions complexity profile plots for four datasets. The same three samples (B1, B2, and D2) in both the HiSeq 2500 and HiSeq 4000 datasets show molecular proportion curves indicative of low library complexity. Indeed even though the number of reads is approximately equal across the 8 samples, the number of molecules differ drastically, by an order of 100 or more, and are lowest in these three samples (see analyses notebooks). Furthermore, whereas the proportion curve of Sample D2 peaks at a moderate PCR duplicate level then tapers off, those of B1 and B2 start picking up and maintain a steady proportion for a wide range of high duplicate level values, up to *r* = 1706 and *r* = 3744, for HiSeq 2500 and HiSeq 4000, respectively. These trends are indicative of problematic samples; incidentally, Samples B1, B2 are the two samples that our analysis identifies to have the largest fraction of phantom molecules. As for the NovaSeq 6000 datasets, the curves for Samples *P* 7_1 and *P* 7_8 indicate low complexity libraries. This is also confirmed by the analysis since both samples turned out to have the two lowest number of molecules but the two highest fractions of phantom molecules among the 16 multiplexed samples.

**Figure 4:**
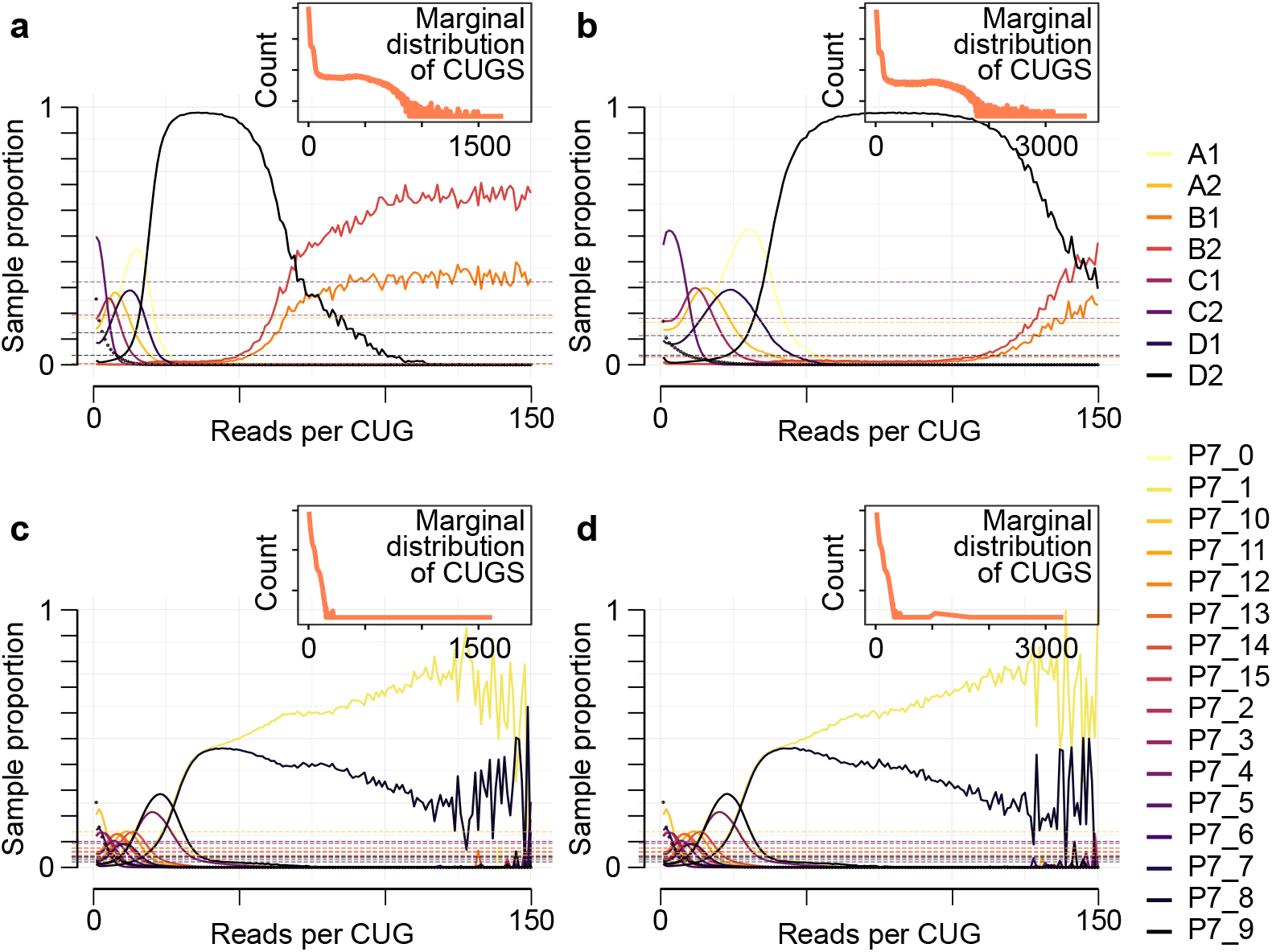
The molecular proportions complexity profile plots for four previously published datasets. Each curve represents the proportion of library complexity (i.e. fraction of molecules) at the indicated PCR amplification level *r* that originate from a given sample. The horizontal dotted lines represent the marginal library complexity proportions. Note that while the plots show *r* up to a maximum of 150, the corresponding subplots in the upper right corners show the distribution of the number of observations for the entire observed range of *r*.

### 6.3 Estimates of the Sample Index Hopping Rate

The model fits for the four datasets are summarized in Table 4 with estimates of *p* along with margin of errors corresponding to the 99 percent confidence intervals. The table also lists the derived estimates of the sample index hopping rates (SIHR) and the sample barcode index hopping rates (SBIHR). As the numbers show, the non-multiplexed (HiSeq 2500) dataset has a much lower (SIHR) than the 3 multiplexed datasets, whose estimates show high consistency. Notice how a slight increase in the sample index hopping rate translates into a sizable increase in the molecule inflation factor and therefore in the proportion of phantom molecules in the dataset.

**Table 4:** Estimates of the sample index hopping rate for one non-multiplexed and three multiplexed sample libraries. 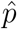 is the estimate of the model parameter (i.e. the complement of the sample index hopping rate, SIHR); SBIHR is the sample barcode index hopping rate (see **Methods** 4.4). *m.e.* is the margin of error corresponding to a 99% confidence interval; *u* is the molecule inflation factor, a summary measure that represents a lower limit on the number of phantom molecules in the data; *g* is the proportion of fugue observations, i.e. CUGs whose reads are all affected by index hopping (and therefore none of their reads represent a real molecule); and PPM=(*u* + *g*)*/*(1 + *u* + *g*) is the Proportion of Phantom Molecules in the dataset. The HiSeq 2500 dataset (control) represents eight mouse epithelial sample libraries that were sequenced separately (non-multiplexed) on different HiSeq 2500 lanes ^[1]^. The HiSeq4000 dataset corresponds to eight mouse epithelial sample libraries that were multiplexed on a single HiSeq 4000 lane ^[4]^. The NovaSeq L1 and NovaSeq L2 datasets correspond to the same set of 16 sample libraries that were multiplexed on two NovaSeq 6000 lanes ^[2]^.

| dataset | $\hat{p}$ | SIHR | SBIHR | $m.e.$ | $u$ | $g$ | PPM |
| --- | --- | --- | --- | --- | --- | --- | --- |
| HiSeq 2500 | 0.99990 | 0.00010 | 0.00011 | $\pm 1.11 \times 10^{-6}$ | 0.00060 | 0.00003 | 0.00062 |
| HiSeq 4000 | 0.99130 | 0.00866 | 0.00960 | $\pm 9.96 \times 10^{-6}$ | 0.08169 | 0.00147 | 0.07688 |
| NovaSeq L1 | 0.99188 | 0.00812 | 0.00853 | $\pm 4.73 \times 10^{-6}$ | 0.04131 | 0.00206 | 0.04167 |
| NovaSeq L2 | 0.99177 | 0.00823 | 0.00864 | $\pm 5.25 \times 10^{-6}$ | 0.04187 | 0.00210 | 0.04222 |

As Fig. 5 shows, the GLM fits the four datasets rather well, especially for low *r* values for which there is almost negligible missingness. Also, notice that the observed chimeric proportions show a downward deviation from the predicted mean trend at high *PCR duplication levels*, especially for those top 1 percent of observations that are characterized by noisiness and sparsity. In fact, the downward trend tends to occur right after the cumulative proportion of molecules in the dataset begins to plateau. The downward trend we observe in the proportion of chimeras seems to indicate a slight decrease in hopping rate that kicks in at high PCR duplication levels. The downward trend in the proportion of chimeras that we observe to deviate from from the model predicted trend could potentially be explained by sample dropout at high *r* values. That is, when the molecules of only a subset of the samples are PCR duplicated at high levels, the multinomial model we specified can no longer assume that we have *S* samples since sample indices cannot hop to nonexistent sample molecules. Accordingly, we expect the curve to differ when we have *S* samples compared to when we have fewer. Another aspect of the observed chimeric proportions becomes apparent at very high *r* values. The observed trend becomes noisy and fluctuating. This can be explained by the the high proportion of missingness at those levels. That is, there are finite observations at a given *PCR duplication level r*, and therefore we will observe only a few of the possible outcomes, especially when *r* > 25, where the number of potential outcomes are in the millions. When this happens, the variability of the mean estimate would increase as the number of observations at a given *r* decreases.

**Figure 5:**
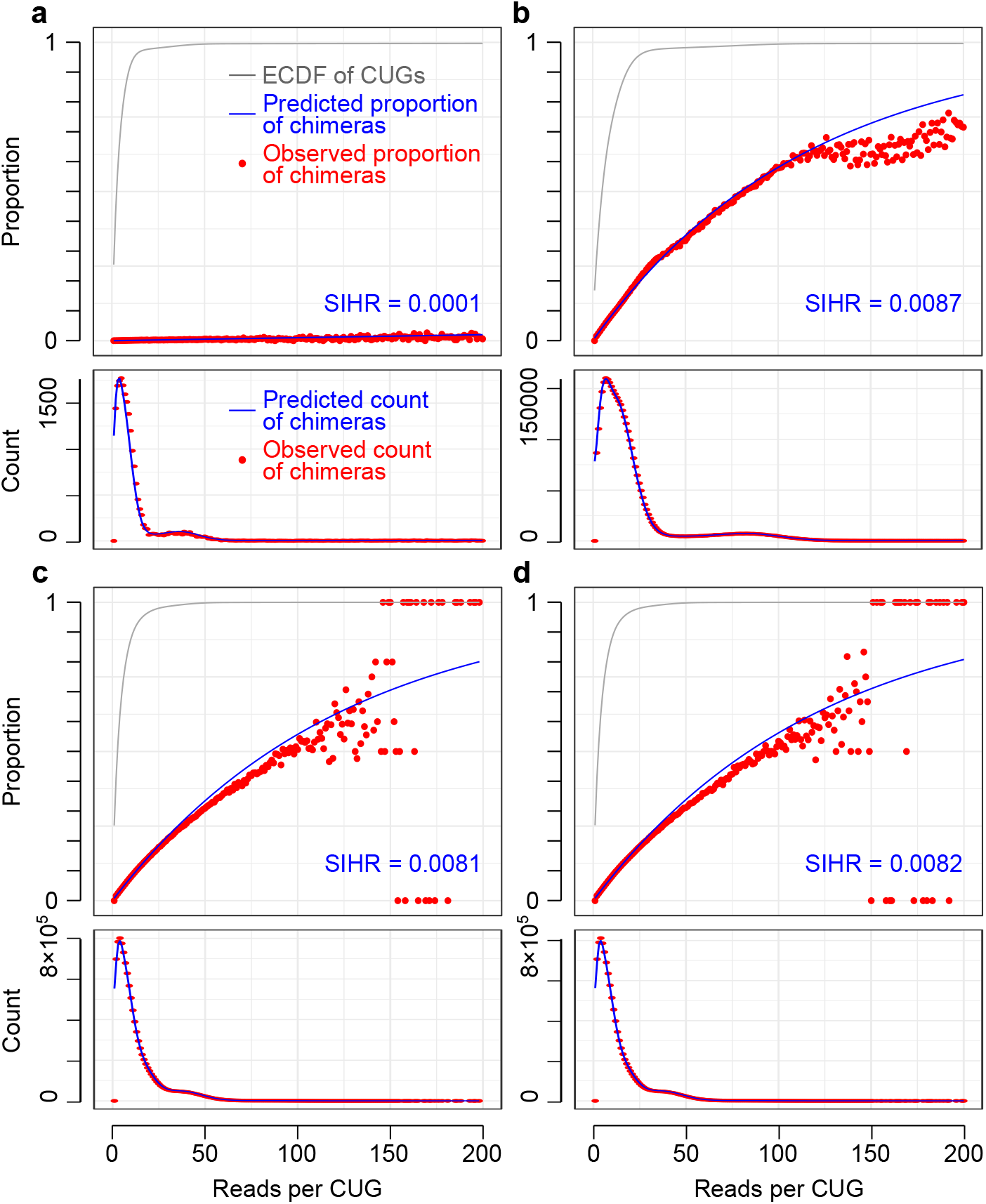
Concordance between model predictions and empirically observed number of chimeras. Each panel corresponds to a different previously published dataset. In each panel, the top plot shows the proportion of chimeras across CUGs with different PCR amplification levels, with red dots depicting observed values and blue line representing the model prediction. The ECDF of *r* (grey) shows the cumulative proportion of observations (CUGs) that are less than or equal to *r*. The bottom plot shows the fit on the absolute count of chimeras rather than proportions. (**a**) Mouse epithelial non-multiplexed HiSeq 2500 dataset^[1]^. (**b**) Mouse epithelial multiplexed HiSeq 4000 dataset^[4]^. (**c**) Tabula Muris NovaSeq 6000 dataset (Lane 1)^[2]^. (**d**) Tabula Muris NovaSeq 6000 dataset (Lane 2)^[2]^.

### 6.4 Minimizing False Positive by Discarding Molecules

As Table 6 shows, leaving the data as it is (i.e. *nopurging*) comes at cost in the form of a large number of false positives (i.e. true phantom molecules that we incorrectly label as real molecules). In contrast, by reassigning reads to the sample with the maximum posterior probability and retaining only those associated molecules (i.e. *nodiscarding*), we are able to substantially reduce the number of false positives at a very little cost of removing true real molecules. We can further minimize the number of false positives by discarding predicted real molecules that have low posterior probability. By using the default *torc* = 3 value, we retain molecules with *tor* values that are equal or greater than the maximum *tor* value not exceeding *torc* = 3. In the table, the rows corresponding to the *above* approach shows the corresponding outcome, inferred sample of origin *s*, posterior probability *qr*, false positive (FP) and false negative (FN) counts, and the *tor* value for the observed outcome with the next higher *tor* value, the one with a lower false positive count.

**Table 5:** Classification performance comparison of four different approaches. Results correspond to a multiplexed dataset of two libraries in which the ground truth is known (i.e. the source sample of each molecule is determined by sequencing the same two libraries on two separate lanes). *No purging* corresponds to the multiplexed data as is. *No discarding* corresponds to reassignment of reads to the sample with the largest posterior probability, but without further filtering. *Discarding* represents the results after discarding the molecules that have a posterior probability lower than a specified cutoff, with the cutoff determined so as to achieve a trade-off ratio of 3 (see Methods). *Min fraction* corresponds to a previous heuristic approach ^[4]^ based on keeping the molecules for which at least 80% of reads are assigned to the same sample. FP: false positive count (phantom molecules that are mistakenly classified as true molecules); FN: false negative count (real molecules that are mistakenly classified as phantom molecules); TP: true positive count (real molecules that are classified as real); TN: true negative count (phantom molecules that are classified as phantom); FPR: false positive rate = FP/(FP+TN); FNR: false negative rate = FN/(FN+TP).

| Approach | FP | FN | TP | TN | FPR | FNR |
| --- | --- | --- | --- | --- | --- | --- |
| No purging | 485399 | 0 | 9717504 | 0 | 1.0000 | 0.0000 |
| Purging at TORC=0 | 12585 | 2590 | 9239510 | 472814 | 0.0259 | 0.0003 |
| Purging at TORC=3 | 10116 | 7685 | 9234416 | 475283 | 0.0208 | 0.0008 |
| Min read fraction $\geq 0.8$ | 10000 | 16094 | 9226006 | 475399 | 0.0206 | 0.0017 |

**Table 6:** Outcomes datatable. The table shows the *tor* cutoff values, corresponding outcome, inferred sample of origin *s*, posterior probability *qr*, false positive (FP) and false negative (FN) counts. Values corresponding to no purging, no discarding, and to the next more conservative *torc* are listed for the mouse epithelial non-multiplexed HiSeq 2500 and multiplexed HiSeq 4000 datasets and the two lanes of the Tabula Muris NovaSeq 6000 datasets.

| dataset | approach | outcome | s | qr | tor | FP | FN |
| --- | --- | --- | --- | --- | --- | --- | --- |
| HiSeq2500 | above | 0,0,1,0,1,0,0,0 | 5 | 0.00630 | 49.64 | 984 | 23,384 |
| HiSeq2500 | torc | 0,0,1,0,0,0,0,0 | 3 | 0.00731 | 2.97 | 1147 | 1201 |
| HiSeq2500 | nodiscarding | 1,0,0,0,0,0,1,0 | 7 | 0.49826 | 0.00 | 1,449 | 303 |
| HiSeq2500 | nopurging |  |  |  |  | 27,393 | 0 |
| HiSeq4000 | above | 0,1,2,1,1,0,0,0 | 3 | 0.10741 | 6.24 | 69,241 | 114,007 |
| HiSeq4000 | torc | 0,0,1,0,0,0,0,0 | 3 | 0.11089 | 2.96 | 80,320 | 25,186 |
| HiSeq4000 | nodiscarding | 0,1,0,0,1,0,1,0 | 5 | 0.56309 | 0.00 | 86,289 | 7,546 |
| HiSeq4000 | nopurging |  |  |  |  | 4,460,074 | 0 |
| NovaSeqL1 | above | 0,0,2,0,0,0,0,0,0,0,2,0,0,0,0,0 | 3 | 0.07375 | 3.01 | 558,115 | 623,953 |
| NovaSeqL1 | torc | 0,0,0,0,0,0,0,0,0,1,1,0,0,0,0,0 | 10 | 0.07399 | 2.99 | 558,550 | 618,509 |
| NovaSeqL1 | nodiscarding | 0,0,0,0,1,0,1,0,1,1,0,0,1,0,0,0 | 10 | 0.72582 | 0.00 | 712,958 | 157,145 |
| NovaSeqL1 | nopurging |  |  |  |  | 11,711,946 | 0 |
| NovaSeqL2 | above | 0,0,1,0,0,0,0,0,0,0,1,0,0,0,1,0 | 3 | 0.07390 | 3.01 | 568,394 | 631,097 |
| NovaSeqL2 | torc | 0,0,0,0,0,0,0,0,0,1,1,0,0,0,0,0 | 10 | 0.07393 | 2.98 | 568,846 | 625,445 |
| NovaSeqL2 | nodiscarding | 0,0,0,1,0,0,0,0,1,1,0,1,0,1,0,1 | 10 | 0.76330 | 0.00 | 725,117 | 159,048 |
| NovaSeqL2 | nopurging |  |  |  |  | 11,902,481 | 0 |

Fig. 6 left shows the trade-off between false negatives and false positives for all four datasets. As can be seen, the optimal cutoff occurs at the point that is closer to the origin. Note that a small marginal decrease in FPs can be offset only by a large increase in FNs. Fig. 7 right shows the marginal increase in false negatives against the marginal decrease in false positives with the default *torc* value in color.

**Figure 6:**
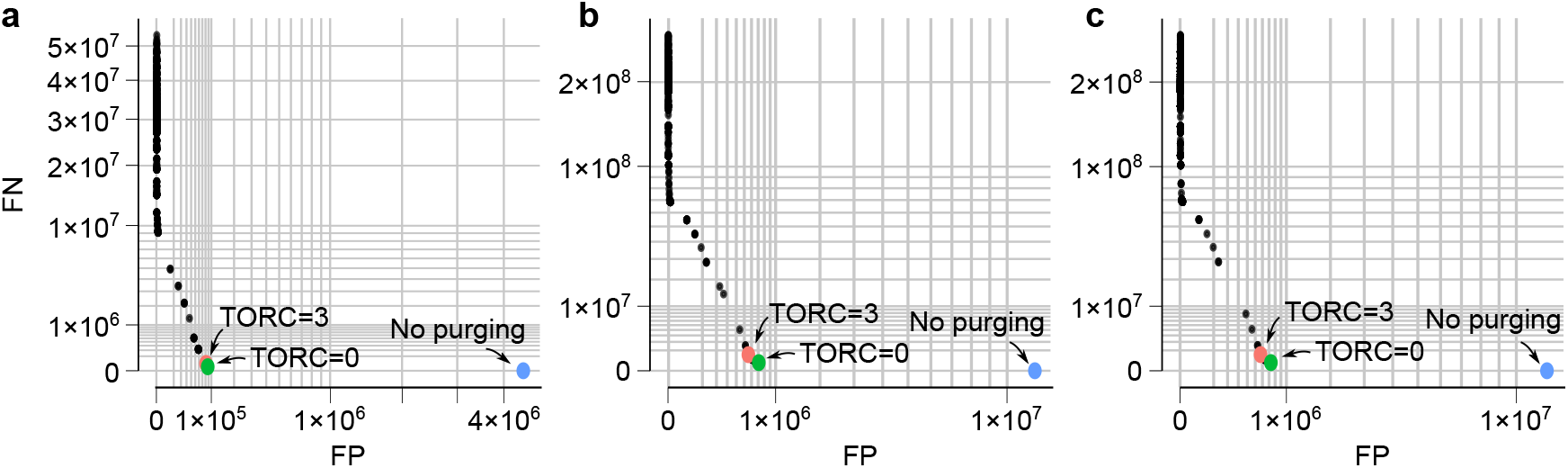
The trade-off between false negatives and false positives. The points correspond to the estimated number of FNs vs. FPs at different posterior probability cutoffs. (**a**) Mouse epithelial multiplexed HiSeq 4000 dataset^[4]^. (**b**) Tabula Muris NovaSeq 6000 dataset (Lane 1)^[2]^. (**c**) Tabula Muris NovaSeq 6000 dataset (Lane 2)^[2]^. In each panel, the FP/FN values when the data are kept as it is (no purging), or reads are reassigned without further filtering by posterior probability (TORC=0) are also shown.

**Figure 7:**
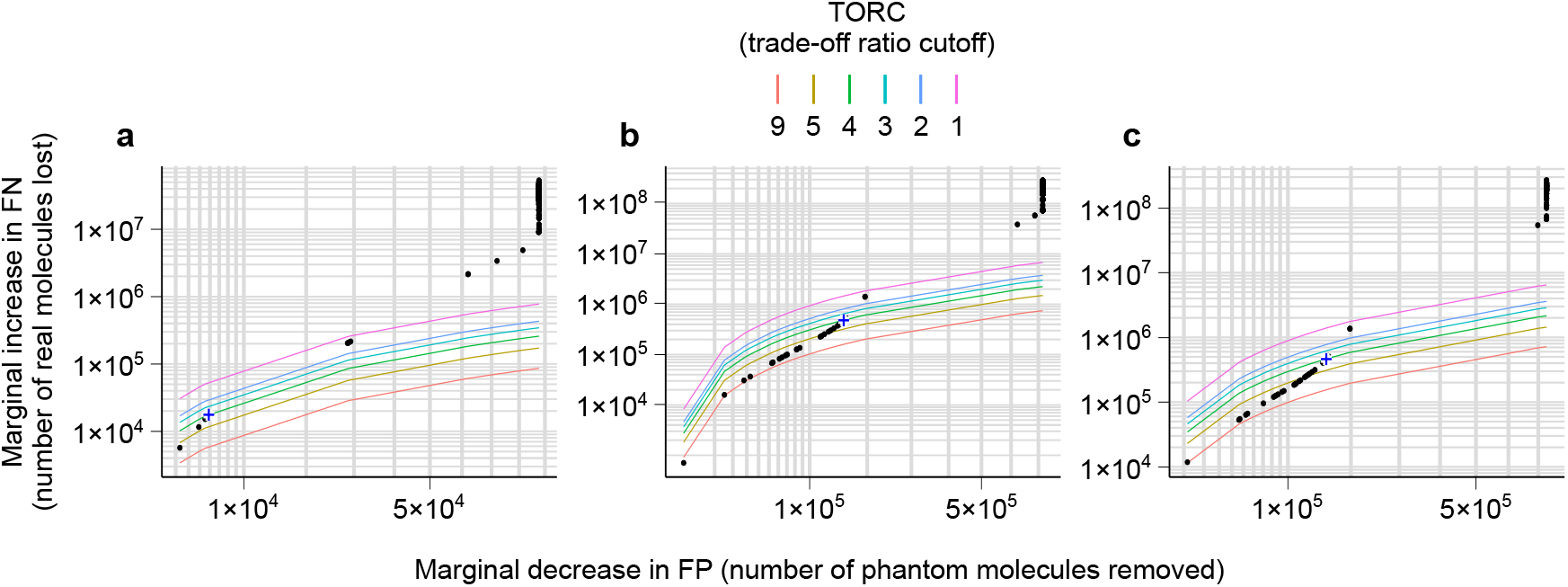
Marginal increase in false negatives against the marginal decrease in false positives. Dots correspond to estimated marginal decrease/increase of false positives/negatives at different posterior probability cutoffs for each dataset. The colored lines show the decision curves for different trade-off ratio cutoff (TORC) values. At any given TORC value, the point that is on or immediately below the corresponding decision curve will be selected as the posterior probability cutoff. (**a**) Mouse epithelial multiplexed HiSeq 4000 dataset^[4]^. (**b**) Tabula Muris NovaSeq 6000 dataset (Lane 1)^[2]^. (**c**) Tabula Muris NovaSeq 6000 dataset (Lane 2)^[2]^.

**Figure 8:**
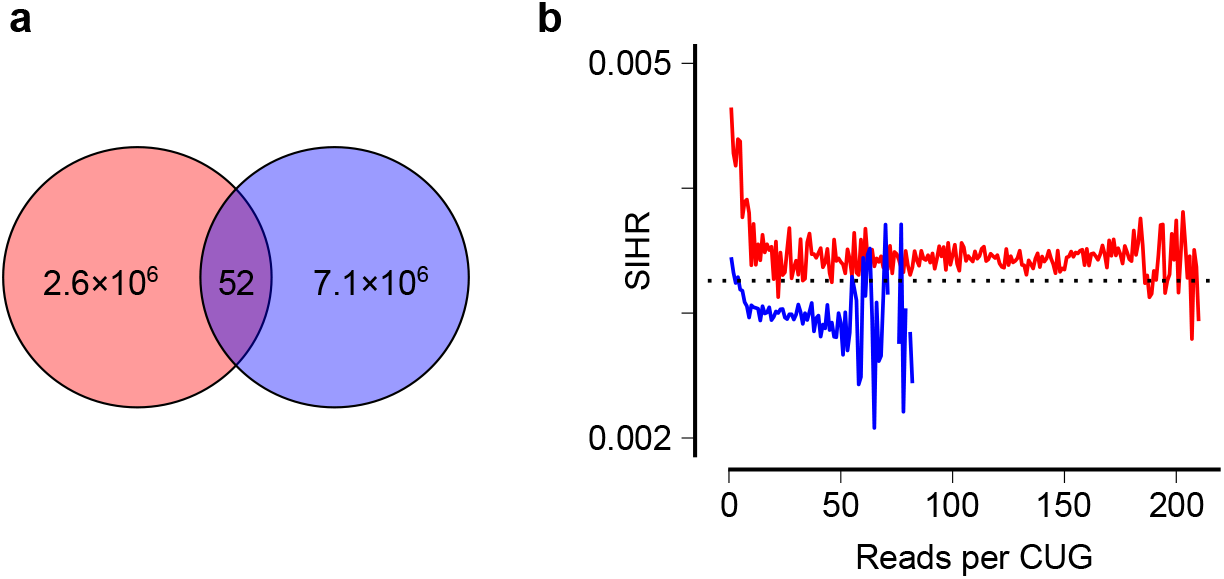
Validating model assumptions. (**a**) Venn diagram showing the number of CUG collisions between two libraries that were sequenced on two separate HiSeq 4000 lanes (i.e. ground truth in the validation dataset, see Methods). Note the minuscule fraction of CUGs that collide (less than 6 in a million), in agreement with Assumption I. (**b**) The proportion of hopped sample indices by source sample across a range of PCR amplification levels *r*, in the multiplexed dataset with known ground truth (the origin of each read was determined based on non-multiplexed sequencing runs of the same libraries on two separate lanes). The straight line shows the marginal mean of proportion of hopped reads. Note that per-sample index hopping rates differ by less than 10%, in agreement with Assumption II. Red: hopping from S1 to S2; Blue: hopping from S2 to S1.

**Figure 9:**
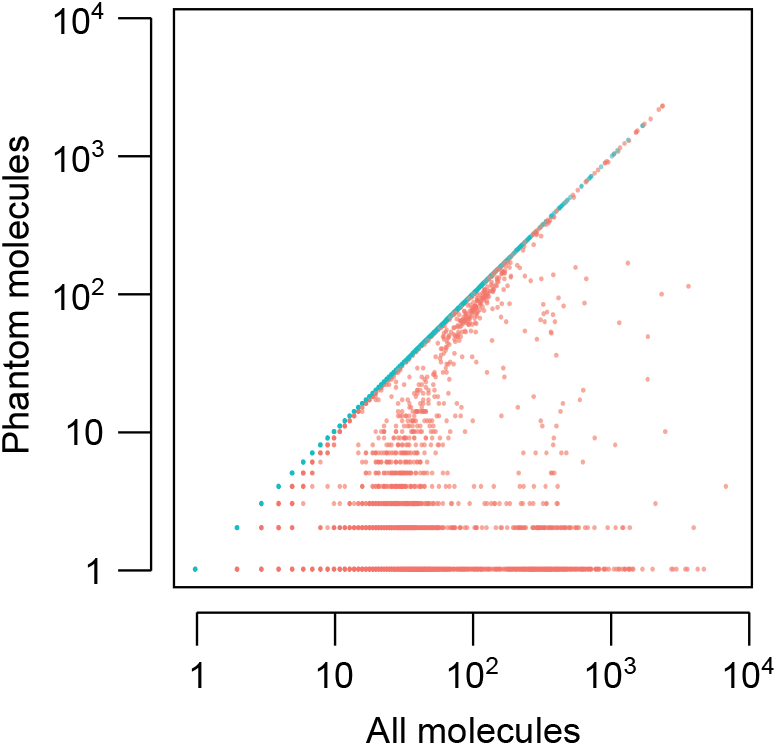
The ratio of phantom to total molecules per cell. Each dot represents one cell that is affected by index hopping. Data correspond to sample S2 from the multiplexed validation dataset (i.e. two libraries with known ground truth multiplexed on the same HiSeq 4000 lane). Blue points are cells whose molecules are all phantom.

**Figure 10:**
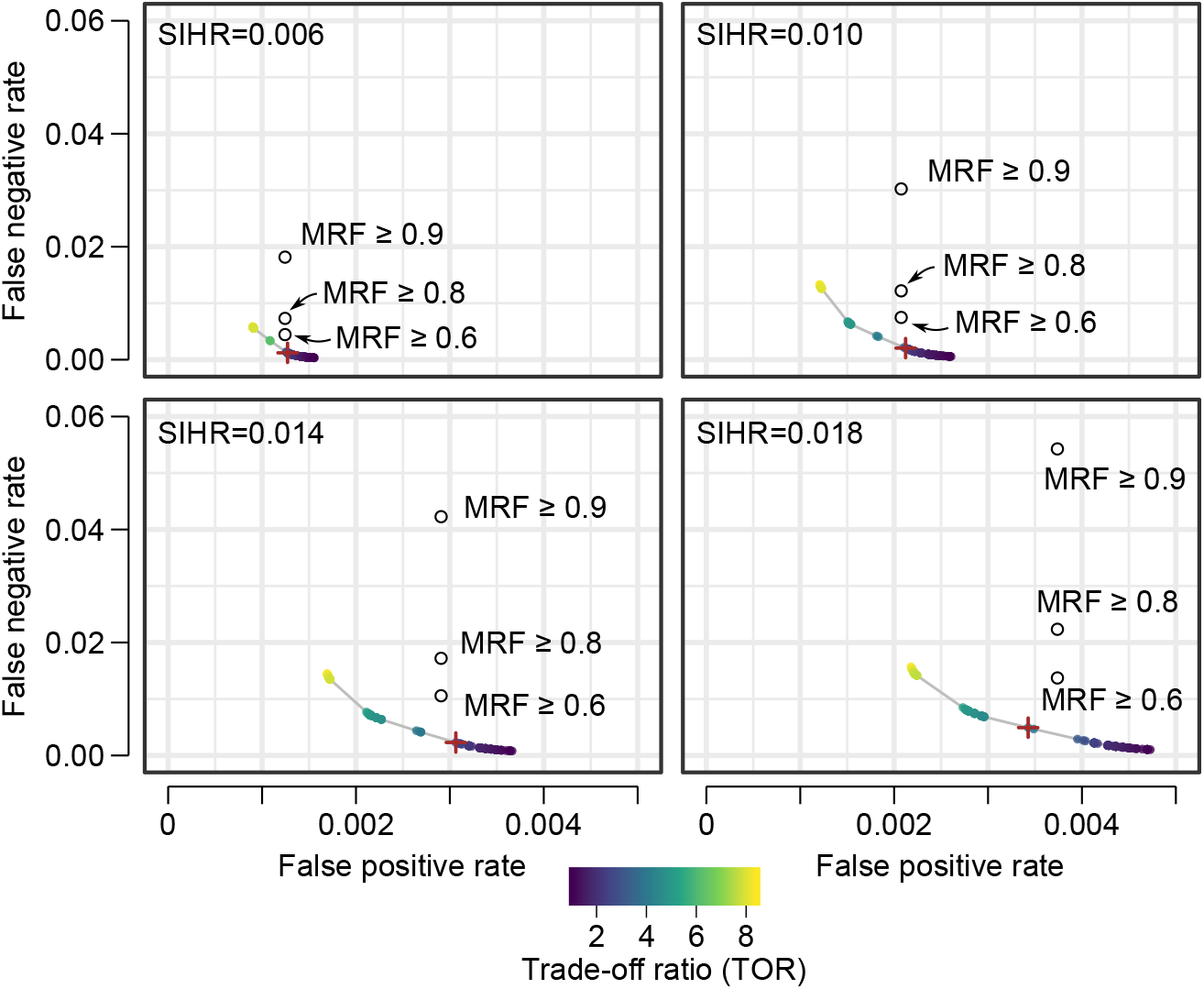
Performance on simulated data. The colored dots represent false positive and false negative proportions after filtering with different posterior probability cutoffs. The color of each dot represents the trade-off ratio, i.e. the number of real molecules lost for every phantom molecule that is correctly purged. The simulations were performed for four evenly spaced values of SIHR (sample index hopping rate). The open circles show the FN/FP values obtained by a previous heuristic approach^[4]^ that is based on retaining CUGs with a certain fraction of reads assigned to only one sample (three choices of the minimum read fraction threshold were used: 0.6, 0.8, and 0.9). The default trade-off ratio cutoff is marked with a plus superimposed on the curve of possible trade-off ratio values within 1-10 range.

### 6.5 The Extent of Contamination by Phantom Molecules

If we had chosen not to decrease the number of false positives by discarding molecules, or in other words if we had set *o*^∗^ = 1, then the number of molecules we predict as phantoms for the entire dataset would be given by (*u* + *g*) × *L*, out of which the number of true phantoms are given by (TN × *L*),and false phantoms by (FN × *L*). However, the proportion of phantoms for individual samples can depart drastically from these marginal measures for the entire dataset as a whole since library complexity varies widely across samples with some samples even consisting mostly of empty cells. Indeed, this is the case for the HiSeq 4000 dataset. As Table 7 shows, the proportion of phantom molecules for this dataset ranges from 0. 012 to 0.864 whereas the same proportion ranges from 0.0132 to 0.176 in both of the two NovaSeq 6000 datasets (see Tables 8 and 8). In contrast, in the HiSeq 2500 dataset, which had not been multiplexed and whose hopping rate we estimated to be extremely small, we see that the range of predicted proportion of phantom molecules ranges from 0.0002 to .02 (Table 7). In the two tables, the number of molecules is given by the second column. The proportion of total molecules predicted as real and therefore retained is given by the *real* column. The proportion of total molecules that predicted as real but discarded since their *q* values fall below the corresponding *tor* cutoff is given by the *false phantoms* column. The proportion of molecules predicted as phantom is given by the *phantoms* column. The two proportions columns give show the discrepancy between library size and library complexity across the samples. The last column measures the extent of PCR Amplification bias as given by *RMR*, the total number of mapped Reads over total number of Molecules (i.e. library size divided by library complexity) ratio. Note that RMR tends to be a good indicator of which samples would have a high proportion of phantoms and also which would have a high proportion of false predicted phantoms (i.e. real molecules which we incorrectly classify as phantoms). For example, Sample *P* 7_8, the sample with the highest RMR in both lanes.

**Table 7:** Extent of contamination by sample for mouse epithelial non-multiplexed HiSeq 2500 dataset (control) and a multiplexed HiSeq 4000 dataset. The column *Molecules* lists the number of total molecules in each sample. The *Breakdown* columns list the proportion of predicted real molecules (*Real*), molecules discarded at TORC=0 (*Phantom*^0^), and additional molecules discarded when TORC is increased to 3 (*Phantom*^3^). The *Proportions* columns list the marginal proportions of molecules and reads across the entire dataset. RMR: the ratio of reads to molecules. Note the small fraction of molecules discarded in the non-multiplexed dataset, which serves as a negative control that should not be affected by our purging approach.

| Sample | Molecules | Breakdown |  |  | Proportions |  |  |
| --- | --- | --- | --- | --- | --- | --- | --- |
|  |  | Real | Phantom<sup>0</sup> | Phantom<sup>3</sup> | Molecules | Reads | RMR |
| HiSeq 2500 Dataset |  |  |  |  |  |  |  |
| A1 | $5.94 \times 10^6$ | 0.9986 | 0.0014 | 0.0000 | 0.13 | 0.13 | 6.1 |
| A2 | $8.02 \times 10^6$ | 0.9997 | 0.0003 | 0.0000 | 0.18 | 0.13 | 4.5 |
| B1 | $0.15 \times 10^6$ | 0.9795 | 0.0205 | 0.0000 | 0.00 | 0.13 | 228.8 |
| B2 | $0.20 \times 10^6$ | 0.9865 | 0.0135 | 0.0000 | 0.00 | 0.13 | 177.8 |
| C1 | $8.51 \times 10^6$ | 0.9997 | 0.0003 | 0.0000 | 0.19 | 0.12 | 3.7 |
| C2 | $14.19 \times 10^6$ | 0.9998 | 0.0002 | 0.0000 | 0.32 | 0.12 | 2.3 |
| D1 | $5.45 \times 10^6$ | 0.9995 | 0.0004 | 0.0001 | 0.12 | 0.11 | 5.5 |
| D2 | $1.63 \times 10^6$ | 0.9984 | 0.0016 | 0.0000 | 0.04 | 0.12 | 20.1 |
| Hiseq 4000 Dataset |  |  |  |  |  |  |  |
| A1 | $7.03 \times 10^6$ | 0.9662 | 0.0337 | 0.0001 | 0.12 | 0.13 | 11.6 |
| A2 | $9.59 \times 10^6$ | 0.9776 | 0.0222 | 0.0002 | 0.17 | 0.13 | 8.6 |
| B1 | $1.82 \times 10^6$ | 0.1351 | 0.8643 | 0.0006 | 0.03 | 0.13 | 44.1 |
| B2 | $1.75 \times 10^6$ | 0.1754 | 0.8239 | 0.0006 | 0.03 | 0.14 | 50.1 |
| C1 | $10.46 \times 10^6$ | 0.9781 | 0.0215 | 0.0004 | 0.18 | 0.12 | 7.3 |
| C2 | $18.65 \times 10^6$ | 0.9874 | 0.0120 | 0.0006 | 0.32 | 0.12 | 4.0 |
| D1 | $6.56 \times 10^6$ | 0.9684 | 0.0314 | 0.0002 | 0.11 | 0.11 | 10.2 |
| D2 | $2.16 \times 10^6$ | 0.8787 | 0.1207 | 0.0006 | 0.04 | 0.12 | 35.1 |

**Table 8:** Extent of contamination by sample for the Tabula Muris NovaSeq 6000 datasets. The column *Molecules* lists the number of total molecules in each sample. The *Breakdown* columns list the proportion of predicted real molecules (*Real*), molecules discarded at TORC=0 (*Phantom*^0^), and additional molecules discarded when TORC is increased to 3 (*Phantom*^3^). The *Proportions* columns list the marginal proportions of molecules and reads across the entire dataset. RMR: the ratio of reads to molecules.

| Sample | Molecules | Breakdown |  |  | Proportions |  |  |
| --- | --- | --- | --- | --- | --- | --- | --- |
|  |  | Real | Phantom<sup>0</sup> | Phantom<sup>3</sup> | Molecules | Reads | RMR |
| NovaSeq 6000 Lane 1 Dataset |  |  |  |  |  |  |  |
| P7_0 | $13.54 \times 10^6$ | 0.9316 | 0.0672 | 0.0012 | 0.05 | 0.07 | 7.9 |
| P7_1 | $8.16 \times 10^6$ | 0.8653 | 0.1343 | 0.0004 | 0.03 | 0.08 | 14.5 |
| P7_2 | $28.02 \times 10^6$ | 0.9754 | 0.0214 | 0.0032 | 0.10 | 0.06 | 2.9 |
| P7_3 | $28.58 \times 10^6$ | 0.9760 | 0.0214 | 0.0026 | 0.10 | 0.06 | 3.1 |
| P7_4 | $8.48 \times 10^6$ | 0.9101 | 0.0898 | 0.0001 | 0.03 | 0.06 | 10.1 |
| P7_5 | $17.40 \times 10^6$ | 0.9678 | 0.0302 | 0.0001 | 0.06 | 0.05 | 4.5 |
| P7_6 | $26.17 \times 10^6$ | 0.9773 | 0.0181 | 0.0045 | 0.09 | 0.04 | 2.3 |
| P7_7 | $11.69 \times 10^6$ | 0.9512 | 0.0478 | 0.0010 | 0.04 | 0.05 | 5.7 |
| P7_8 | $5.99 \times 10^6$ | 0.8247 | 0.1753 | 0.0000 | 0.02 | 0.07 | 17.3 |
| P7_9 | $10.49 \times 10^6$ | 0.9146 | 0.0849 | 0.0005 | 0.04 | 0.07 | 10.1 |
| P7_10 | $38.68 \times 10^6$ | 0.9837 | 0.0132 | 0.0031 | 0.14 | 0.06 | 2.4 |
| P7_11 | $16.20 \times 10^6$ | 0.9549 | 0.0437 | 0.0014 | 0.06 | 0.07 | 6.1 |
| P7_12 | $21.28 \times 10^6$ | 0.9663 | 0.0315 | 0.0022 | 0.08 | 0.07 | 4.8 |
| P7_13 | $20.80 \times 10^6$ | 0.9681 | 0.0298 | 0.0021 | 0.07 | 0.07 | 4.9 |
| P7_14 | $12.92 \times 10^6$ | 0.9470 | 0.0523 | 0.0007 | 0.05 | 0.06 | 6.8 |
| P7_15 | $12.81 \times 10^6$ | 0.9600 | 0.0386 | 0.0014 | 0.05 | 0.05 | 5.4 |
| NovaSeq 6000 Lane 2 Dataset |  |  |  |  |  |  |  |
| P7_0 | $13.59 \times 10^6$ | 0.9309 | 0.0012 | 0.0679 | 0.05 | 0.07 | 7.9 |
| P7_1 | $8.20 \times 10^6$ | 0.8645 | 0.1351 | 0.0004 | 0.03 | 0.08 | 14.4 |
| P7_2 | $28.11 \times 10^6$ | 0.9750 | 0.0218 | 0.0032 | 0.10 | 0.06 | 2.9 |
| P7_3 | $28.67 \times 10^6$ | 0.9756 | 0.0217 | 0.0026 | 0.10 | 0.06 | 3.1 |
| P7_4 | $8.51 \times 10^6$ | 0.9088 | 0.0910 | 0.0001 | 0.03 | 0.06 | 10.1 |
| P7_5 | $17.43 \times 10^6$ | 0.9674 | 0.0306 | 0.0020 | 0.06 | 0.05 | 4.5 |
| P7_6 | $26.25 \times 10^6$ | 0.9770 | 0.0184 | 0.0046 | 0.09 | 0.04 | 2.3 |
| P7_7 | $11.72 \times 10^6$ | 0.9506 | 0.0485 | 0.0010 | 0.04 | 0.05 | 5.7 |
| P7_8 | $6.03 \times 10^6$ | 0.8238 | 0.1762 | 0.0000 | 0.02 | 0.07 | 17.3 |
| P7_9 | $10.53 \times 10^6$ | 0.9139 | 0.0856 | 0.0005 | 0.04 | 0.07 | 10.1 |
| P7_10 | $38.78 \times 10^6$ | 0.9834 | 0.0135 | 0.0031 | 0.14 | 0.06 | 2.4 |
| P7_11 | $16.25 \times 10^6$ | 0.9543 | 0.0443 | 0.0014 | 0.06 | 0.07 | 6.1 |
| P7_12 | $21.34 \times 10^6$ | 0.9657 | 0.0320 | 0.0022 | 0.08 | 0.07 | 4.8 |
| P7_13 | $20.85 \times 10^6$ | 0.9676 | 0.0303 | 0.0022 | 0.07 | 0.07 | 4.9 |
| P7_14 | $12.96 \times 10^6$ | 0.9463 | 0.0530 | 0.0007 | 0.05 | 0.06 | 6.8 |
| P7_15 | $12.84 \times 10^6$ | 0.9595 | 0.0392 | 0.0014 | 0.05 | 0.05 | 5.4 |

**Table 9:** Effect of contamination on cell calling in mouse epithelial non-multiplexed HiSeq 2500 (control) and multiplexed HiSeq 4000 datasets. The *cell* and *empty* columns list the number of cell-barcodes that were categorized as RNA-containing cells or background cells (empty droplets), respectively. The *Phantom* column enumerates the cells and empty droplets that disappear once phantom molecules are purged; The *Consensus* column enumerates the cells and empty droplets that maintain the same status no matter whether the phantom molecules were purged or not; The *Reclassified* column represents the number of cell-barcodes that were re-classified (transitioned) as empty droplet or cell after purging the phantom molecules. Note the small number of changes in the non-multiplexed dataset, which serves as a negative control that should not be affected by our purging approach.

| Sample | Phantom |  | Consensus |  | Reclassified |  |
| --- | --- | --- | --- | --- | --- | --- |
|  | Cell | Empty | Cell | Empty | Cell | Empty |
| **HiSeq 2500 Dataset** |  |  |  |  |  |  |
| A1 | 0 | 1,332 | 409 | 123,769 | 1 | 0 |
| A2 | 0 | 770 | 564 | 138,948 | 0 | 2 |
| B1 | 4 | 1,258 | 199 | 26,138 | 0 | 17 |
| B2 | 0 | 1,136 | 16 | 29,971 | 0 | 0 |
| C1 | 0 | 808 | 739 | 145,283 | 0 | 0 |
| C2 | 0 | 668 | 1,316 | 195,614 | 4 | 1 |
| D1 | 0 | 778 | 685 | 98,969 | 1 | 0 |
| D2 | 0 | 975 | 160 | 61,353 | 1 | 0 |
| **HiSeq 4000 Dataset** |  |  |  |  |  |  |
| A1 | 0 | 12,478 | 401 | 138,117 | 13 | 105 |
| A2 | 0 | 11,188 | 565 | 156,913 | 13 | 105 |
| B1 | 0 | 70,193 | 16 | 40,016 | 0 | 1,007 |
| B2 | 0 | 65,001 | 20 | 45,503 | 0 | 885 |
| C1 | 0 | 11,729 | 730 | 168,503 | 0 | 118 |
| C2 | 0 | 10,087 | 1,318 | 232,404 | 5 | 154 |
| D1 | 0 | 11,609 | 661 | 119,473 | 15 | 64 |
| D2 | 0 | 15,912 | 165 | 82,287 | 7 | 29 |

### 6.6 The Effects of Phantom Molecules on Identifying RNA-containing Cells

Given that a substantial percentage of mapped molecules in the dataset originate from empty droplets, one can argue that what matters in the final analysis is whether there turns out to be a substantial degree of phantom contamination in the subset of molecules originating from RNA-containing droplets. In general, there are two potential effects that phantom molecules bring about: (1) the emergence of phantom cells and more commonly (2) causing cell-calling algorithms to miss-classify empty droplets as cells and vice versa.

To assess the extent and effects of contamination by phantom molecules in this subset, we used the *EmptyDrops* ^[8]^ algorithm to call cell-barcodes in order to determine whether a cell-barcode originated from a cell or an empty droplet. For each dataset, we ran *EmptyDrops* the first time on the unpurged data and then a second time of purged data. To compare the results of the two runs, all barcodes classified as either cell or empty by the algorithm were divided into three groups each as shown in Tables 9 and 10. In the tables, the *Consensus* columns show the number of cell-barcodes that maintain the same classification as cell or empty, across the two runs. The *Phantom* columns show the number of cell-barcodes whose associated molecules were all predicted as phantom molecules (true and false phantoms). The *Re-classified* columns show the number of barcodes that have been reclassified after running the cell-calling algorithm of purged data. For example, we see that for the A1 sample in the HiSeq 4000 dataset, 13 barcode that was previously classified as background, were re-classified as cells after purging whereas 105 cell-barcodes that were classified as cell before purging were re-classified as empty after purging. For sample, the extent of the reclassification is more pronounced. If we had called cells without first purging the dataset of phantom molecules, we would have believed that we have called a total of 1023 (= 16+1007) cells, whereas re-running cell-calling on purged data would have produced no more than 16 cells.

**Table 10:** Effect of contamination on cell calling in the Tabula Muris NovaSeq 6000 datasets. The *cell* and *empty* columns list the number of cell-barcodes that were categorized as RNA-containing cells or background cells (empty droplets), respectively. The *Phantom* column enumerates the cells and empty droplets that disappear once phantom molecules are purged; The *Consensus* column enumerates the cells and empty droplets that maintain the same status no matter whether the phantom molecules were purged or not; The *Reclassified* column represents the number of cell-barcodes that were re-classified (transitioned) as empty droplet or cell after purging the phantom molecules.

| Sample | Phantom |  | Consensus |  | Reclassified |  |
| --- | --- | --- | --- | --- | --- | --- |
|  | Cell | Empty | Cell | Empty | cell | empty |
| **NovaSeq 6000 Lane 1 Dataset** |  |  |  |  |  |  |
| P7_0 | 0 | 61,772 | 1,119 | 245,922 | 156 | 196 |
| P7_1 | 0 | 73,677 | 551 | 231,820 | 105 | 223 |
| P7_2 | 0 | 53,833 | 4,037 | 306,452 | 13 | 21 |
| P7_3 | 0 | 49,661 | 3,974 | 330,196 | 26 | 26 |
| P7_4 | 0 | 63,917 | 927 | 250,569 | 29 | 69 |
| P7_5 | 0 | 47,393 | 19,69 | 316,514 | 37 | 12 |
| P7_6 | 0 | 45,701 | 7,268 | 304,202 | 63 | 17 |
| P7_7 | 0 | 63,518 | 733 | 210,349 | 21 | 11 |
| P7_8 | 0 | 95,414 | 10,89 | 175,093 | 21 | 108 |
| P7_9 | 0 | 76,342 | 2,306 | 238,232 | 8 | 89 |
| P7_10 | 0 | 41,213 | 2,449 | 375,818 | 19 | 11 |
| P7_11 | 0 | 64,228 | 3,474 | 259,465 | 1 | 32 |
| P7_12 | 0 | 59,249 | 3,027 | 291,810 | 16 | 28 |
| P7_13 | 0 | 53,450 | 3,383 | 305,486 | 10 | 26 |
| P7_14 | 0 | 64,676 | 2,601 | 244,136 | 12 | 28 |
| P7_15 | 0 | 54,118 | 2,753 | 242,666 | 12 | 8 |
| **NovaSeq 6000 Lane 2 Dataset** |  |  |  |  |  |  |
| P7_0 | 0 | 62,320 | 947 | 246,383 | 358 | 213 |
| P7_1 | 0 | 73,684 | 611 | 232,303 | 43 | 224 |
| P7_2 | 0 | 53,673 | 4,036 | 307,442 | 11 | 22 |
| P7_3 | 0 | 50,035 | 3,981 | 331,060 | 24 | 25 |
| P7_4 | 0 | 64,176 | 826 | 251,665 | 17 | 173 |
| P7_5 | 0 | 47,130 | 2,023 | 317,367 | 40 | 18 |
| P7_6 | 0 | 46,122 | 7,264 | 305,496 | 76 | 16 |
| P7_7 | 0 | 63,731 | 732 | 211,401 | 23 | 11 |
| P7_8 | 0 | 95,323 | 1,089 | 176,709 | 24 | 120 |
| P7_9 | 0 | 76,811 | 2,305 | 239,238 | 13 | 95 |
| P7_10 | 0 | 41,633 | 2,451 | 376,946 | 21 | 13 |
| P7_11 | 0 | 64,495 | 3,476 | 261,000 | 0 | 38 |
| P7_12 | 0 | 59,618 | 3,029 | 293,207 | 16 | 33 |
| P7_13 | 0 | 54,118 | 3,389 | 306,711 | 7 | 27 |
| P7_14 | 0 | 65,386 | 2,597 | 245,290 | 15 | 33 |
| P7_15 | 0 | 54,121 | 2,757 | 244,201 | 6 | 8 |

For the NovaSeq 6000 datasets, the effects of contamination are less severe than those see in the HiSeq 4000 dataset. For many samples the number of barcodes that are reclassified as empty can make a substantial fraction of total cells (e.g. Sample *P* 7_01 with 223 barcodes). Note that we opted to treat the two lanes separately for the cell-calling stage of the analysis to evaluate the consistency of the proposed approach on two datasets that only differ by flowcell sequencing lane. For downstream analysis, data from the two lanes should be combined before cell-calling is performed.

### 6.7 Validation

#### Assumption I

Corollary 1 of Assumption 1 states that the probability of a CUG collision is zero. To examine the extent of the assumption’s validity, we counted the number of colliding CUGs in the inner joined data table. We found that there are only 52 (real, real) colliding CUGs out of a total of 9,252,147, or in other words, a collision rate of approximately 0.00001 for the case of two samples. Accordingly, for practical purposes, we can consider Assumption I to be valid given that collision rate is close to zero and is more than 3 orders of magnitude smaller than the CUG collision rates observed in data contaminated by sample index hopping.

#### Assumption II

Assumption 2 states that the hopping probability of a sequencing read’s sample index is the same regardless of either the sample index’s source or target. To validate the assumption we counted the fraction of hopped reads originating from each sample at the entire range of PCR duplication levels (see Fig. 8). As the figure shows, the estimated sample index hopping rate 0.00320±0.00001 is very close to the marginal ground truth estimate of 0.00326 (i.e. the true mean). That said there are two trends that seem to slightly depart from the model’s assumption. First, there is a minor difference between the two samples’ proportion of hopped reads, *SIHR*_1_ = 0.00346 vs *SIHR*_2_ = 0.00302 (the subscript denotes the source sample). Second, the ground truth estimates for both samples start out at higher proportion values but stabilize starting at *r* = 10 (see the reproducible R markdown notebook *validation_hiseq4000_2.nb.html* for details). These trends could be sub-sampling artifacts of the filtering procedure step in which we only retained CUGs that are observed in both the multiplexed and non-multiplexed samples (each of which has twice the number of reads). Alternatively, these trends could be persistent across experiments in which case the model we have proposed captures rather well but not perfectly the underlying mechanism of sample index hopping. A model that is governed by several sample-specific hopping rates that are also dependent on the number of duplicated reads would provide better accuracy but at the cost of intractable computational and mathematical complexity. Even then, the improved accuracy in estimating the hopping rate would not affect the purging procedure greatly given that the molecular complexity profiles plays an important role in assigning reads to their sample of origin.

#### 6.7.1 Comparing and Evaluating Classification Performance

We evaluate the tor cutoff approach by contrasting it with two alternative actions: no purging, where we leave the data as it is, and no discarding, where we purge predicted phantom molecules but refrain from discarding real molecules whose inferred sample-of-origins have low posterior probabilities. We also compare the classification performance of the proposed method to the minimum read fraction molecule exclusion approach^[4]^, in which the sample-of-origin molecule is deemed the one with the majority of the reads (default ≥ 80%).

As Table 5 and Fig. 1, the *torc* approach achieves a false positive count 10,116 that is very close to the lowest possible (i.e. 9,995, the number of fugue molecules in the dataset) while maintaining a low number of false negatives, 7,685 as well. In contrast, although the maximum fraction approach achieves a lower number of false positives, the reduction of 116 false positives is offset by a substantial number of false negatives, 16,094, an increase of 8,409 over *torc* approach.

#### 6.7.2 Extent of contamination

Although the proportion of hopped reads in Sample 1 and Sample 2 are 0.0035 and 0.0030, respectively, the corresponding proportion of phantom molecules (PPM) are an order of magnitude higher, at 0.085 and 0.037, respectively. Furthermore, the percentage of cells that contain at least one phantom cell is approximately 20%. For illustration see Fig. 9 which plots, for each affected cell, the number of phantom molecules against the total number of molecules.

### 6.8 Simulation

To compare the performance of the proposed approach to the minimum read fraction molecule exclusion approach^[4]^ for a case when there are more than two samples, as was the case in the validation data, we simulated data for (*S* = 8) samples using the molecular complexity profile computed for the mouse epithelial cells HiSeq 4000 data. For computational feasibility, outcomes were simulated for all *r* values up to a maximum of 15 for a range of four *p* values. The simulation results show (see Table 11 and Fig. 10) that in contrast to the minimum read fraction approach, the tor (default maximum value not exceeding 3) approach gives a false negative proportion that is about 3 times lower while also maintaining a comparable false positive proportion, which progressively becomes relatively smaller as the hopping rate increases. More importantly, as the tor curves show in Figure 10, the entire range of possible tor cutoff values provide a more optimal trade-off choice than the minimum read fraction approach (represented by three possible thresholds *mf* = (0.6, 0.8, 0.9)) for balancing the false negative and positive rates.

**Table 11:**
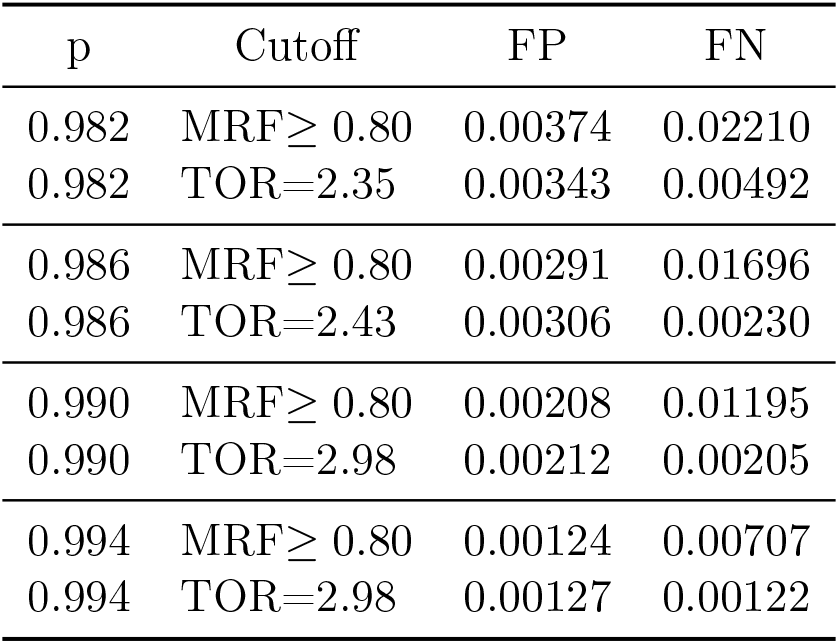
Performance on simulated data. The table shows the false positive and false negative proportions for the default cutoff value of the minimum read fraction (MRF) method^[4]^ and our probabilistic modeling approach for four evenly spaced values of *p* (the complement of the sample index hopping rate). We used a trade-off ratio cutoff (TORC) of 3 for our method. Note that the trade-off ratio (TOR) after purging corresponds to the largest TOR value obtainable that is still smaller than TORC=3.

## Acknowledgements

The authors thank Alfredo Staffa and James Webber for their comments and useful discussions, and Lorenzo Ferri and Veena Sangwan for providing the validation samples.

## Authors’ Contributions

Methodology and conceptualization: RF and HSN. Formal analysis, mathematical derivation, and code implementation: RF. Visualization: RF and HSN. Writing (original draft): RF and HSN. Writing (review and editing): RF, HSN, JR, HD. Validation Data Acquisition: JR and HD. Directing the study and funding acquisition: HSN.

## Supplementary Note

### 1 Mathematical Derivations

#### 1.1 The Molecular Proportions Complexity Profile

Let 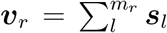 denote the molecule count across the *S* samples at level *r* and 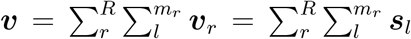 the molecule count across the *S* samples for the entire data. Since E(***v***_*r*_) = *m*_*r*_ × ***π***_*r*_, the expected total molecule count is therefore 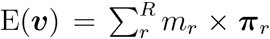. If ***π***_*r*_ = ***π*** for all *r* = 1,…,*R*, then E(***v***) = *L* × ***π*** correspond to the *library complexities* of the *S* samples.

Let 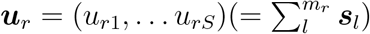 represent the unobserved molecule (i.e. UMIs) counts at *PCR duplication level r* across *S* samples. The distribution of ***u***_*r*_ can be obtained by noting that sum of *m*_*r*_ identically distributed categorical random variables with parameter vector ***π***_*r*_ is a multinomial random variable with the same parameter vector. That is,

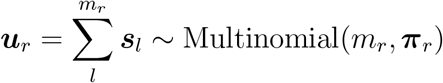

Now, the expected molecule count across samples in the entire data is given as such.

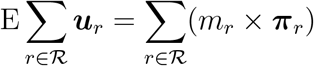

However, we do not observe ***s***_*l*_, and the expected total number of observed molecules in the data can be much greater due to the presence of phantom molecules brought about by sample index hopping.

Let 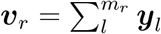 denote the total read count at *PCR duplication level r*. In what follows, it helps to think of the *m*_*r*_ observations as members of *S* partitions, where the size of partition *s* is given by *π*_*rs*_, the proportion of molecules in Sample *s* at *PCR duplication level r*. Now since

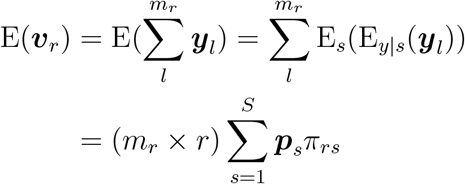

therefore

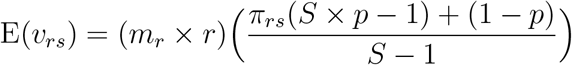

and so we can obtain an estimate for *π*_*rs*_ from the observed proportion of read counts 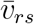 for sample *s* observations at *PCR duplication level r*. That is,

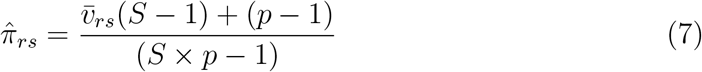

Notice that when there is no sample index hopping (i.e. *p* = 1), the proportion of molecules equals the proportion of reads, that is 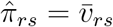. Also, note that for the denominator to be positive, the proportion of reads 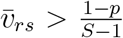 for all *s* = 1, …, *S*. In empirical data, when this relationship is violated, we can set 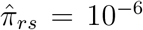 and renormalize 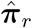 accordingly.

#### 1.2 The Distribution of k-chimeras

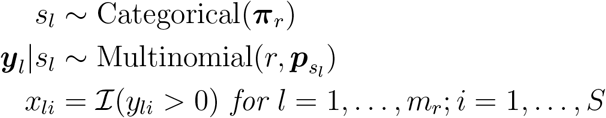

where 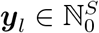, **p** ∈ [0, 1]^*S*^, ||***p***||_1_ = 1, and *S* ∈ ℕ

The marginal distribution of each element of the multinomial observation ***y***_*l*_ is binomial. That is,

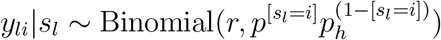

where the Iverson bracket notation is used. It follows that the distribution of *x*_*li*_ is a Bernoulli with a probability parameter that can be obtained by observing that the expectation of an event defined by an indicator function is equal to the probability of that event. That is,

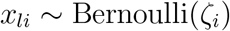

where

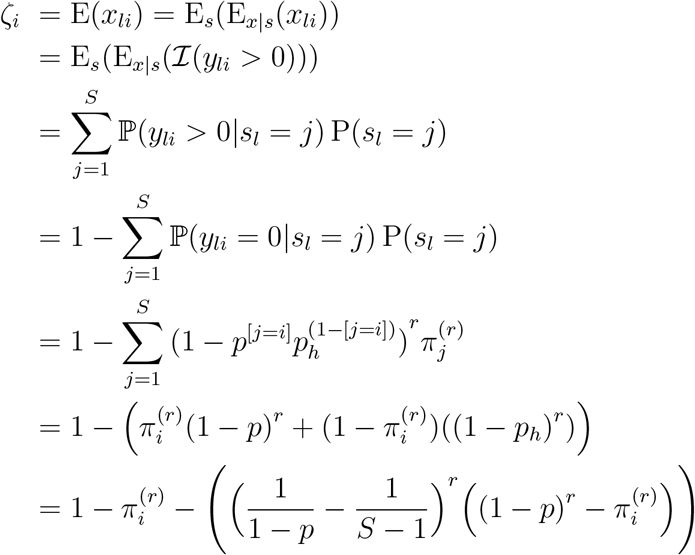

Now given that the Bernoulli random variables (i.e. the elements of ***x***_*r*_) can be treated as independent (for *r* > *S*) but not identically distributed, their sum, which indicates the category of the observation (i.e. *k*–chimera), can be given by the Poisson Binomial distribution. That is,

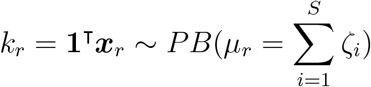

and a PMF given by the following recursive formula ***ζ***

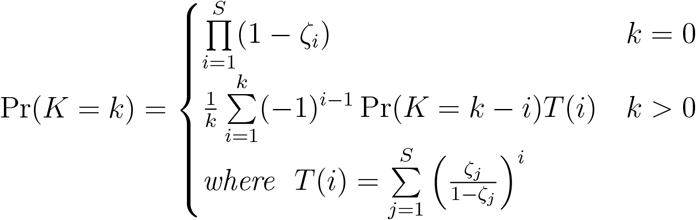

Note the independence assumption does not hold when *r* < *S* since Pr(*K* = 0) is not zero even though it should be given that the *r* reads must be belong to at least one sample. Also note that the mean, *µ*_*r*_, of the Poisson Binomial distribution is equal to the sum of the probabilities. In this case, the 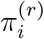’s do not affect *µ*_*r*_ since they cancel out. That said, they do affect the variance 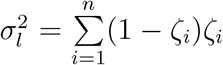 where it is maximized when the *ζ*_*i*_’s, and accordingly the 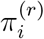’s, are equal. In this case *k*_*l*_ can be reduced to the sum of dependent Bernoulli variables, which for *r* > *S* can be approximated as the sum of independent and identically distributed Bernoulli variables. That is,

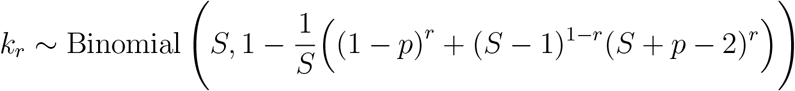

The distribution of phantoms can be obtained from *k*_*r*_ by noting that for non-fugue observations, the number of phantoms equals *k*_*r*_ − 1. The expected fraction of phantoms at *PCR duplication level r* is therefore approximately as such.

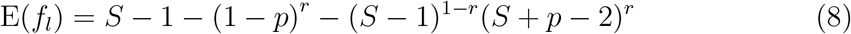

Figure 1 plots the functional relationship between the expected fraction of phantoms and the variables *p*, *r*, and *S*.

**Figure 1:**
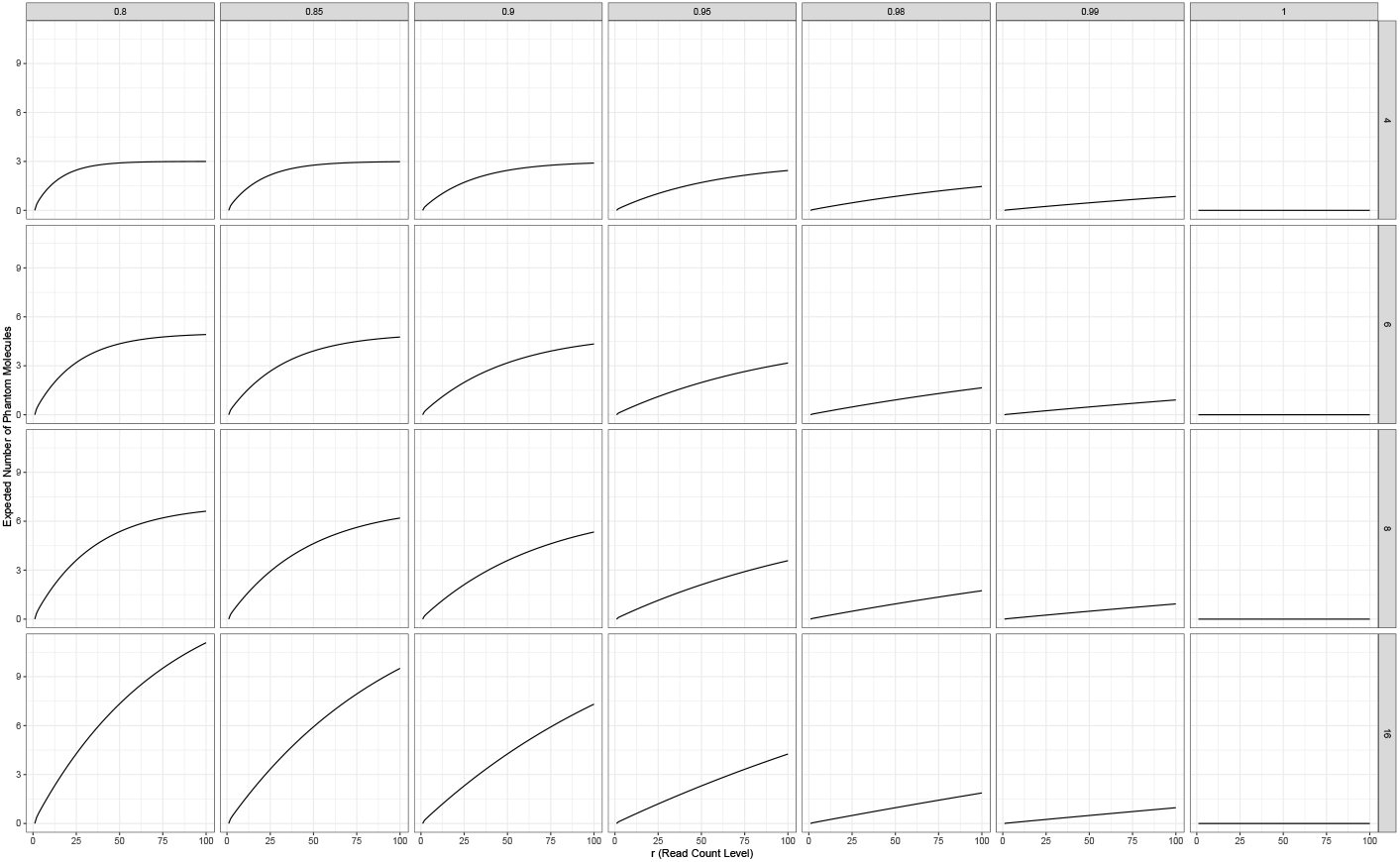
The expected number of phantoms for an observation with *PCR duplication level r* for a range of values of *p* and number of samples *S*

#### 1.3 Distribution of non-chimeras

For the case of *k* = 1, or non-chimeric observations, we can derive a closed form of the distribution by noting that a *non-chimera* is a count observation ***y***_*l*_ for which (*y*_*li*_ = *r*) for any *i* ∈ {1, …, *S*}. We denote the event of observing a *non-chimera* by a Bernoulli random variable 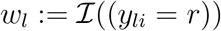 with parameter *p*_*w*_ given by

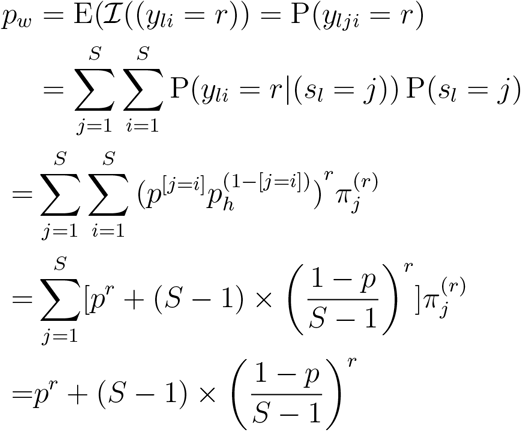

As a result, the distribution of observing a *non-chimera* is given by

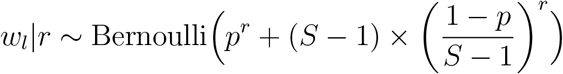

where 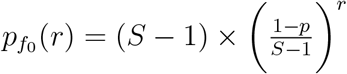 is the probability of observing a non-chimeric fugue observation with *r* reads and *p*^*r*^ is the probability of observing a non-chimeric non-fugue observation with *r* reads.

#### 1.4 Estimation of the Sample Index Hopping Rate

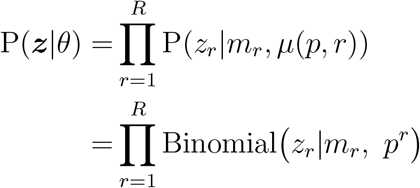

Note that the mean function *µ*(*r*) = *p*^*r*^ can be expressed as such.

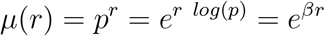

which corresponds to the integral curve solution of the differential equation

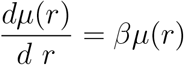

This is a negative growth curve since *β* = *log*(*p*) ∈ (−∞, 0). We can estimate *β* by formulating the problem as a generalized linear regression for binomial counts with a systematic component *η* = *βr* and a log link function *g*(*µ*) = *log*(*µ*) such that.

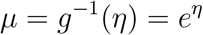

Note that the standard link function for binomial counts is the 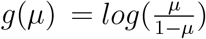. However, it corresponds to the solution of another differential equation, namely.

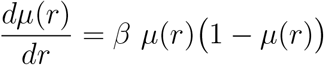

Although the log link function is not symmetric, *η* is defined only on the negative real line. Furthermore, the *log* link renders the parameters interpretable since *β* is basically just the log of *p* the parameter of interest.

The relationship between the number of non-chimeras *z*_*r*_ and the sample index hopping rate (1 − *p*) at a given *PCR duplication level r* and with an *m*_*r*_ number of observations can be formulated as a *generalized linear regression* model with a *log* link function as follows.

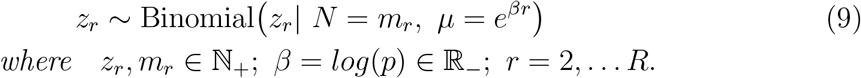

An estimate of the sample index hopping rate can be obtained from the regression coefficient as such.

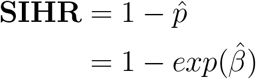

We can use the estimate of the sample index hopping rate to obtain the expected number of hopped reads at each *PCR duplication level*, namely (*m*_*r*_ × *r* × **SIHR**). We can also plug it into Equation 8 to compute the expected Fraction of Phantom Molecules (**FPM**) for the entire dataset. That is.

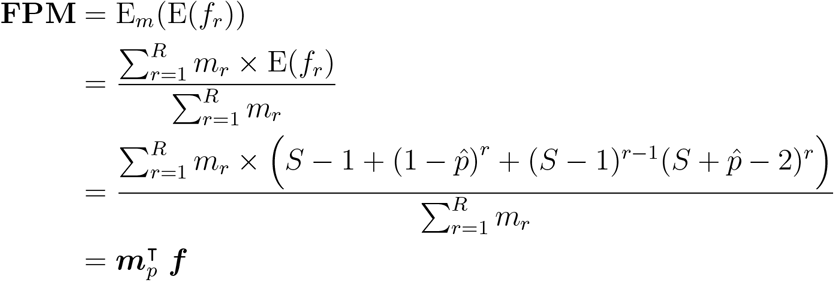

Here 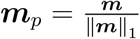 where ***m*** = (*m*_1_,…, *m*_*R*_) and ***f*** = (E(*f*_1_,…,E(*f*_*R*_)).

We would like to note that we can do away with Assumption 3 and estimate *p* in Model 4 using a likelihood optimization procedure. However, the optimization algorithm might not converge or it can get stuck at a local optimum. Furthermore, compared to the GLM approach, it is not straightforward to obtain standard errors or any measure of uncertainty by optimizing the likelihood. We provide code in the paper’s GitHub repository showing the likelihood optimization approach and its overall similar results to the GLM approach.

#### 1.5 Estimating the Sample Barcode Index Hopping Rate

A sample in a 10X Genomics experiment is actually labelled by, not one, but four different barcodes. By considering only the sample labels instead of the sample barcode, the statistical approach outlined in this paper models sample index hopping across samples rather than across individual barcodes. Nonetheless, we can still estimate the sample barcode index hopping rate by noting that when we treat all reads of a given sample as having the same sample index, we are effectively collapsing a 4 × *S*-dimensional multinomial vector of barcode sample index hopping probabilities ***p***_*b*_ into an *S*-dimensional vector of sample index hopping probabilities ***p***. That is, since the read counts are modeled with a multinomial, we can fuse the four barcode count events within a sample by summing their read counts and corresponding probabilities. Accordingly, the relationship between the two parameters of interest *p*_*b*_ and *p* can be expressed as such.

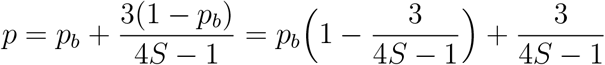

where *p*_*b*_ is probability that a given sample barcode index does not hop. Accordingly, we can obtain an estimate of the sample barcode index hopping rate by expressing *p*_*b*_ in terms of *p* and *S*.

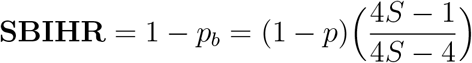

Notice that the multiplicative factor is at its maximum (i.e. 1.75) when *S* = 2. As *S* increases however, it approaches 1 and the two terms become approximately equal.

### 1.6 The Posterior Distribution of the True Sample of Origin

The posterior distribution can be obtained as follows.

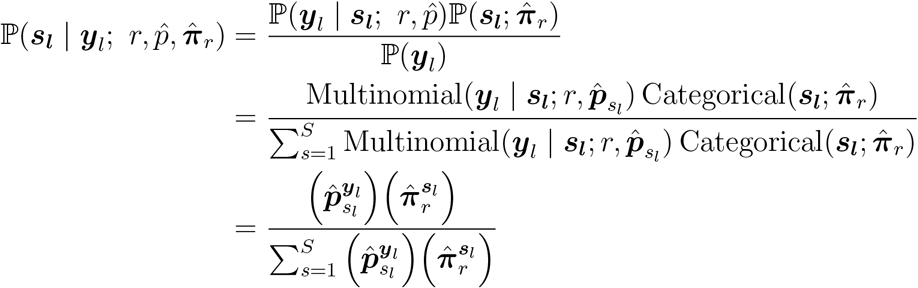

which for each element, simplifies to

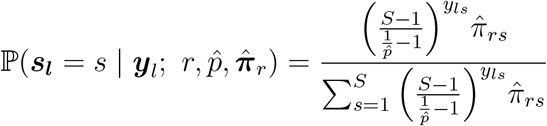

#### 1.7 Youden’s J statistic

The cut-off given by *Youden’s* **J** is optimal in the sense that it minimizes the probability of random guessing when we give equal weight to FPR and FNR, or equivalently to sensitivity and specificity^[13]^. Nonetheless, optimizing **J** might not be the most appropriate choice for the problem at hand since the number of phantom molecules is an order of magnitude smaller than the number of real molecules. That is, FPR and FNR are proportions of unbalanced classes, whose denominators depend on *u*, which is a function of the hopping rate, and *g*, which not only depends on the hopping rate, but whose estimate can be biased.

#### 1.8 Computing the Proportion of Fugue Observations (g)

To obtain the proportion of all fugues **g**, we need to take into account the proportion of chimeric fugues **g**_**c**_ as well the proportion of non-chimeric fugues, which we denote by **g**_**0**_. We can compute **g**_**0**_ by multiplying the observed relative frequency proportion vector of the read count levels (i.e. 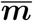) by the vector of non-chimeric fugue probabilities, whose elements are given by 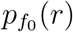.

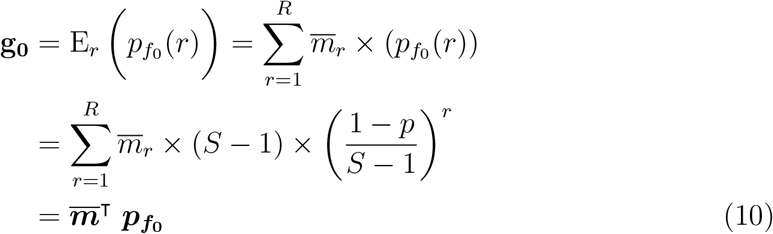

The probability of observing chimeric fugues can be obtained as such.

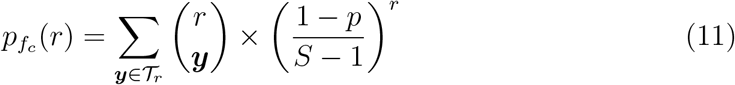

where the set 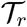 consists of outcomes corresponding to events where all the *r* reads hopped to two or more target samples: the outcomes (0, 2, 1) and (0, 1, 2) in Figure 3. For *r* = 1, the set 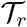 is empty; for *r* = 2, there are 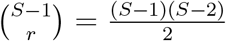 such outcomes. For *r* > 3, it becomes increasing hard to enumerate and compute the probabilities for all the possible outcomes, especially when *S* is large as well.

However, the value of 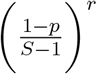 becomes negligibly small for *r* > 3 given that *p* tends to be very close to 1. Furthermore, empirical data is characterized by a highly skewed distribution of *PCR duplication levels* such that the proportion of observations in the first few *r* values makes up a substantial fraction of all observations. Therefore even if 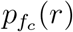 was not negligible for *r* > 3, it would ultimately be weighted down by the respective 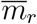. Under this assumption, the proportion of fugue observations can therefore be accurately approximated as such.

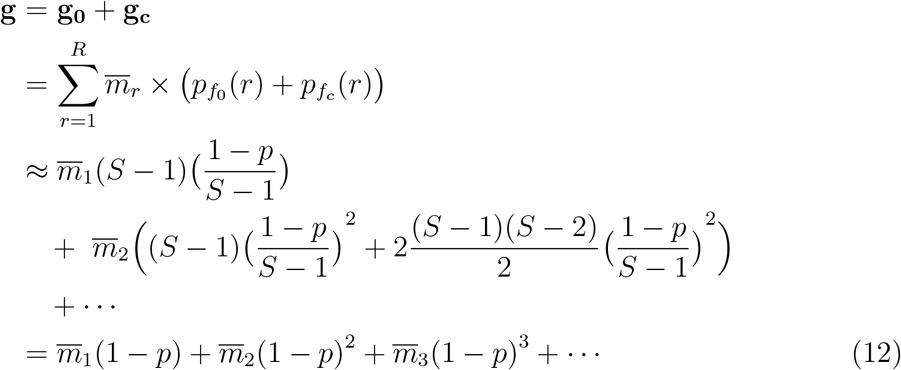

For example when *p* = .99, 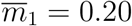, 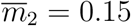, and 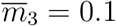 **g** = 0.0020151. Or in other words, only 0.2 % of observations are fugues. If all the data is restricted to have *PCR duplication level r* = 1 (i.e. 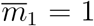), then **g** = (1 − *p*) would be the maximum possible proportion of fugue observations.

### 2 Overview of Computational Workflow

Here we provide a rough summary of the data analysis steps that comprise the *PhantomPurge* workflow. The code implementing the workflow and a set of reproducible R Markdown notebooks detailing the data analysis steps are available on the paper’s GitHub repository.

#### Step 1 creating a joined read counts table

In order to create the joined data table of gene-mapped read counts across samples, we load from the *molecule_info.h5* file of each sample the data corresponding to the following four fields: *cell-barcode*, *gene*, *umi*, and *reads*. We then join the data from all samples into a single data table that is keyed by a unique *cell barcode-gene-umi* combination ID. In addition to the unique label ID, each row of the data table contains read counts across all the *S* samples.

#### Step 2 creating an outcome counts table

Given that the number of observations in the joined read counts data table tends to be in the hundreds of millions, we then collapse observations with similar outcomes and tally their frequency of occurrence. Working with a data table of outcome tallies drastically reduces the number of computations required. A few grouping variables are also added to the joined data table of outcome tallies to be used for subsequent analysis steps. The resulting data table contains the following additional variables: an outcome character variable (e.g. "(1,0,1,1)") to group the observations into unique outcomes, a count variable (*n*) denoting the number of times a particular outcome was observed in the data, a *PCR duplication level* variable (*r*), and a (*k_chimera*) variable that counts the number of molecules we observe in each outcome.

#### Step 3 estimating the sample index hopping rate

Using the introduced grouping variable, we tallied the number of observations that are chimeric and those that are non-chimeric at each *PCR duplication level r*. The resulting data table is then used as input to the GLM Model 5 in order to estimate, from the proportion of chimeric observations, the sample index hopping rate.

#### Step 4 inferring the true sample of origin and reassigning reads

To compute the posterior probability of the true sample index (the latent variable *s*) for each unique observed outcome, we first summarized from the data the conditional and marginal distributions of reads. The summaries are represented by a vector of *PCR duplication level* proportions 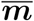, which we used to compute the empirical marginal distribution of *q*, and a set of *I* across-samples read count proportion vectors from which we estimated the proportion of molecules across samples 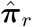, one at each *PCR duplication level r*, that in turn we plugged in to calculate the conditional posterior probability *q*|***y***. For each outcome, the posterior probabilities of the possible true samples of origin are computed and the index of the sample with the maximum posterior probability along with posterior probability itself is added to the original joined read count table. We then used the predicted true sample of origin and its associated posterior probability to reassign reads to their predicted sample of origin.

#### Step 5 determining the cutoff for purging predicted phantom molecules

In order to remove predicted phantom molecules from the data while minimizing the rate of false positives and false negatives, we computed the trade-off ratio (tor) statistic by dividing the marginal increase in FNs over the marginal decrease of FPs for each observed unique *qr*\* value. Predicted real molecules associated with outcomes whose corresponding tor values are greater than the user-specified *torc* are retained (default value is 3). To purge the data, the read counts are first deduplicated to obtain a table of molecule (i.e. UMI) counts. After purging, the molecule counts are collapsed over gene labels to produce a gene-by-cell umi-count expression matrices for all the samples sequenced in the same lane.

#### Step 6 identifying RNA-containing cells before and after purging

To distinguish cells from empty droplets, the cell-calling algorithm *EmptyDrops* ^[8]^ is called. The set of called cells in each sample is used to filter out background cell-barcodes from the deduplicated purged UMI count matrices. The filtered matrices are then saved to file in order to be used for downstream statistical analyses. By comparing the results of the cell-calling algorithm from both purged and unpurged data, one can then examine the extent of contamination of phantom molecules on data that would have otherwise not been purged.

### 3 Method’s Limitations

The work presented in this paper has the following limitations.

#### The non-negligible probability of a *cell-umi-gene* label collision across samples when the number of samples is large

As the number of multiplexed samples increases, Corollary 1 would no longer hold since the probability of observing, in more than one sample, a given *cell-umi-gene* label combination becomes non-negligible. That said, in single cell experiments, multiplexing more than 16 samples on a single lane is not commonly done given the smaller library size and lower genomic coverage that follows as a result. Furthermore, the adoption of a longer UMI index in the latest 10X Genomics Single Cell 3’ v3 assay would further reduce the probability of potential collisions, thus rendering this concern less of a problem.

#### The sensitivity of the FPR-minimizing cutoff on the ECDF of *q*

As we have already mentioned, the marginal distribution of *q* given by Equation 6 has no closed-form nor is it feasible to compute the exact probabilities for all its possible outcomes even when *r* is not large. As *r* increases, we tend to observe in the data an increasingly fewer proportion of all possible outcomes and subsequently we would expect the resulting ECDF to deviate from the theoretical distribution. That said, the deviations would only affect the classification measures and the determination of the cutoff *o*^∗^ whether optimally or not, but they would not affect the read reassignment procedure and the initial purging that retains all predicted real molecules (i.e. when we set *o*^∗^ = 1) since the marginal distribution is not involved at this step.

#### Memory requirements

For data generated on the Hiseq 4000, the analysis workflow does not require computational requirements beyond what is found on a regular desktop with 32G of RAM. However, for Novaseq 6000 data with a large number of multiplexed samples (e.g. *S* > 8), more memory would be needed - up to 150G if not more. In particular, the first step in the workflow consisting of joining data from all samples into a datatable keyed by cell-barcode, UMI, and gene ID is the most memory intensive.

#### Incorporating the sample barcode indices in the model

In this work, we formulated a model for index hopping that treats all the four distinct sample barcodes present in a given sample as identical. Even though we showed in Section 1.5 that the sample barcode index hopping rate can be derived from the model nonetheless, we have not used information from the sample barcode indices to reassign reads or to purge phantom molecules. The model can potentially be expanded to incorporate the 4 sample barcodes, but such an extension would be accomplished at the cost of increased, if not prohibitive, computational and memory requirements since the workflow would need to start with the FASTQ files, not the output of the CellRanger pipeline and furthermore, the read count data table would need to be joined and keyed across 4 times the number of samples, potentially requiring more memory than would be available on some clusters. We would like to point out that the main reason we chose to formulate a model for index hopping at the level of samples and not individual sample barcodes is that in the end, sample index hopping between reads belonging to the same sample would not have any effect on downstream analyses since no phantom molecules would be generated when a read in a given sample swaps its sample barcode index with one of the other three barcode indices that are assigned to the same sample.

## Footnotes

1 As measured by the total number of nucleotide sequence reads generated in a single run of an experiment

